# Whole-genome sequencing of 128 camels across Asia provides insights into origin and migration of domestic Bactrian camels

**DOI:** 10.1101/656231

**Authors:** Liang Ming, Liyun Yuan, Li Yi, Guohui Ding, Surong Hasi, Gangliang Chen, Tuyatsetseg Jambl, Nemat Hedayat-Evright, Mijiddorj Batmunkh, Garyaeva Khongr Badmaevna, Tudeviin Gan-Erdene, TS Batskh, Wenbin Zhang, Azhati Zulipikaer, Hosblig, Erdemt, Arkady Natyrov, Prmanshayev Mamay, Narenbatu, Gendalai Meng, Choijilsuren Narangerel, Orgodol Khongorzul, Jing He, Le Hai, Weili Lin, Sirendalai, Sarentuya, Aiyisi, Yixue Li, Zhen Wang, Jirimutu

## Abstract

The domestic Bactrian camels were treated as the principal means of locomotion between the eastern and western cultures in history. To address the question of their origin, we performed whole-genome sequencing of 128 camels across Asia, including representative populations of domestic Bactrian camels from the Mongolian Plateau to the Caspian Sea, as well as the extant wild Bactrian camels and dromedaries. The domestic and wild Bactrian camels showed remarkable genetic divergence since they were split from dromedaries, confirming they were separated species. The wild Bactrian camels made also little contribution to the ancestry of domestic ones. Among the domestic Bactrian camels, those from Iran exhibited the largest genetic distance from others, and were the first population to separate in the phylogeny. Although evident admixture was observed between domestic Bactrian camels and dromedaries living around the Caspian Sea, the large genetic distance and basal position of Iranian Bactrian camels could not be explained by introgression alone. Taken together, our study favored the Iranian origin of domestic Bactrian camels, which were then immigrated eastward to Mongolia where the native wild Bactrian camels inhabited. This study illustrated the complex genomic landscape of migration underlying domestication in Bactrian camels.

## INTRODUCTION

Camels (*Camelus, Camelini*) contain two extant domestic species, the one-humped dromedary (*Camelus dromedarius*) and the two-humped Bactrian camel (*Camelus bactrianus*) [1, 2]. While the former herds are mainly feed in North Africa and West Asia, the latter herds wildly live in the cold desert areas of Northeast and Middle Asia. The wild Bactrian camel (*Camelus ferus*), the only representative of the wild tribe Camelini as a result of the extinction of the wild dromedary [3], is listed as critically endangered by the International Union for Conservation of Nature [4] with an estimation of a few hundreds to 2,000 individuals [5, 6]. Historically, the wild Bactrian camel was widely distributed throughout Asia, extending from the great bend of the Yellow River westward to central Kazakhstan, but it can only be found in the Mongolian Gobi and the Chinese Taklimakan and Lop Noor deserts today [7]. Fossil and molecular evidence suggested that the ancestor of camels lived in North America and spread to Asia via the Bering land bridge around 11 or 16 million years ago [8, 9]. Within the Camelini, the dromedaries and Bactrian camels were then split around 4 or 5 million years ago [9, 10]. The domestication of camels, like many other domestic mammals, has promoted unprecedented progress in cultural and economic development of human societies, representing a great leap forward for human civilization. For example, the Bactrian camels were rightfully considered as the principal means of locomotion across the bridge between the Eastern and Western cultures in the time of the Silk Road.

The origin of domestic dromedaries was recently revealed by world-wide sequencing of modern and ancient mitochondrial DNA (mtDNA), which suggested that they were at first domesticated in the southeast Arabian Peninsula [11]. However, the origin of domestic Bactrian camels is still a mystery. One intuitive possibility was the extant wild Bactrian camels were the progenitor of the domestic form, which were then dispersed from the Mongolian Plateau to west gradually [7, 12]. This hypothesis was supported by the presence of Camelid faunal remains at Neolithic sites near Mongolia, although it was unclear these were the domestic as opposed to the wild ones [12]. Despite the fact that the wild Bactrian camel has certain characters distinct from the domestic one, such as the lower, pyramid-shaped humps, the thinner, lithe legs, and the smaller and more slender body [7], it is still difficult to tell them apart based on their morphological features alone. Nevertheless, molecular studies based on mtDNAs discovered dramatic sequence variations between the wild camels and domestic Bactrian camels, and proposed that the extant wild Bactrian camel was a separate lineage and not the direct progenitor of the domestic Bactrian camel [9, 13]. Another possible place of origin was Iran [1], where early skeletal remains of domestic Bactrain camels (around 2,500-3,000 BC) were discovered [14]. Although prehistoric mtDNAs of Bactrian camels supported the idea that the domestication took place in the southwestern Middle Asia rather than in Mongolia or East Asia [15], the incomplete archaeological findings and limited molecular markers provided little decisive information about the actual domestication history.

Whole-genome sequences contain much more molecular markers than mtDNAs, which were successfully used to portray the origin, migration and admixture of humans [16-18] and domestic animals [19] such as dogs [20-23] and pigs [24-26]. The genome assembly of Bactrian camels were published recently [10, 27], which provided a new opportunity to examine the evolutionary relationship between the extant wild and domestic Bactrian camels and trace their origin. In this study, we performed whole-genome sequencing of 128 camels including both domestic and extant wild Bactrian camels from their typical habitats. We included dromedaries as well because they were not only the outgroup of Bactrian camels in phylogeny, but also had a long history of hybridization with Bactrian camels in breeding practice [2, 28], especially in Middle Asia. Our results supported the Iranian-origin hypothesis of domestic Bactrian camels, and a roughly west-to-east route of migration back to the Mongolian Plateau.

## RESULTS

### Sample collection and whole genome sequencing

A total of 105 domestic Bactrian camels across Asia, 19 wild Bactrian camels from Gobi-Altai region in Mongolia as well as 4 dromedaries from Iran were gathered for this study (Supplementary Figure S1 and Table S1). The domestic Bactrian camels were chosen to cover as many major geographic regions as possible, including 55 from Inner Mongolia (IMG), Xinjiang (XJ) and Qinghai (QH) of China, 28 from Mongolia (MG), 6 from Kazakhstan (KAZA), 10 from Russia (RUS) and 6 from Iran (IRAN). As a variety of local breeds were formed due to the wide utilization of domestic Bactrian camels in China and Mongolia, 8 different representative breeds were chosen from the regions. The other domestic Bactrian camels from Middle Asia were living around the Caspian Sea.

After DNA extraction, individual genomes were sequenced to an average of 13× coverage (Supplementary Figure S2 and Table S2). The sequence reads were aligned to our previous genome assembly of the Bactrian camel [27] for variant calling (Supplementary Figure S3). After stringent filtering, we totally identified 15.76 million SNPs and 1.97 million small indels (Supplementary Table S3). Functional annotation of the variants indicated that about 68.75% of them were intergenic, 30.18% were intronic and 0.77% were exonic (Supplementary Table S4 and Table S5). Among the variants, 15.49 million, 5.96 million and 10.31 million were identified in the domestic Bactrian camels, wild Bactrian camels and dromedaries, respectively (Supplementary Table S3). Although dromedaries were more divergent from both of the Bactrian camel species in phylogeny, the domestic Bactrian camels shared more variants with the dromedaries (52.69%) than with the wild Bactrian camel (31.14%) (Supplementary Figure S4) due to the tremendous reduction of genetic variants observed in the extant wild Bactrian camel.

### Genetic diversity and differentiation

For a more detailed comparison of the genetic diversity among different populations, we first removed 14 individuals showing close genetic relationship with the remaining others (Supplementary Table S6). The pairwise nucleotide diversity π (Figure 1A) of dromedaries (1.65× 10^−3^) was significantly higher than that of Bactrian camels from all geographic regions (0.92×10^−3^-1.14×10^−3^, Supplementary Table S7), which was in contrast to previous heterozygosity estimates based on individual genomes [10]. One important reason could be the hybridization practice between dromedaries and Bactrian camels in Middle Asia [28]. Among the Bactrian camels, the wild population showed the lowest π (0.92×10^−3^) compared with all of the domestic populations (Figure 1A and Supplementary Table S7). Although this result violated many cases that wild animals usually have higher genetic diversity than the domestic breeds derived from them such as dogs [22], pigs [25] and rabbits [29], it would happen due to the fact that the extant wild Bactrian camels are critically endangered with an extremely small population size [4]. In addition, the domestic populations living around the Caspian Sea generally showed a higher diversity than those living in the Mongolia Plateau (Figure 1A). The tendency was also confirmed by the Watterson’s θ (Supplementary Figure S5). Again, the hybridization with dromedaries in Middle Asia could account for the higher diversity in the region.

**Figure 1.**
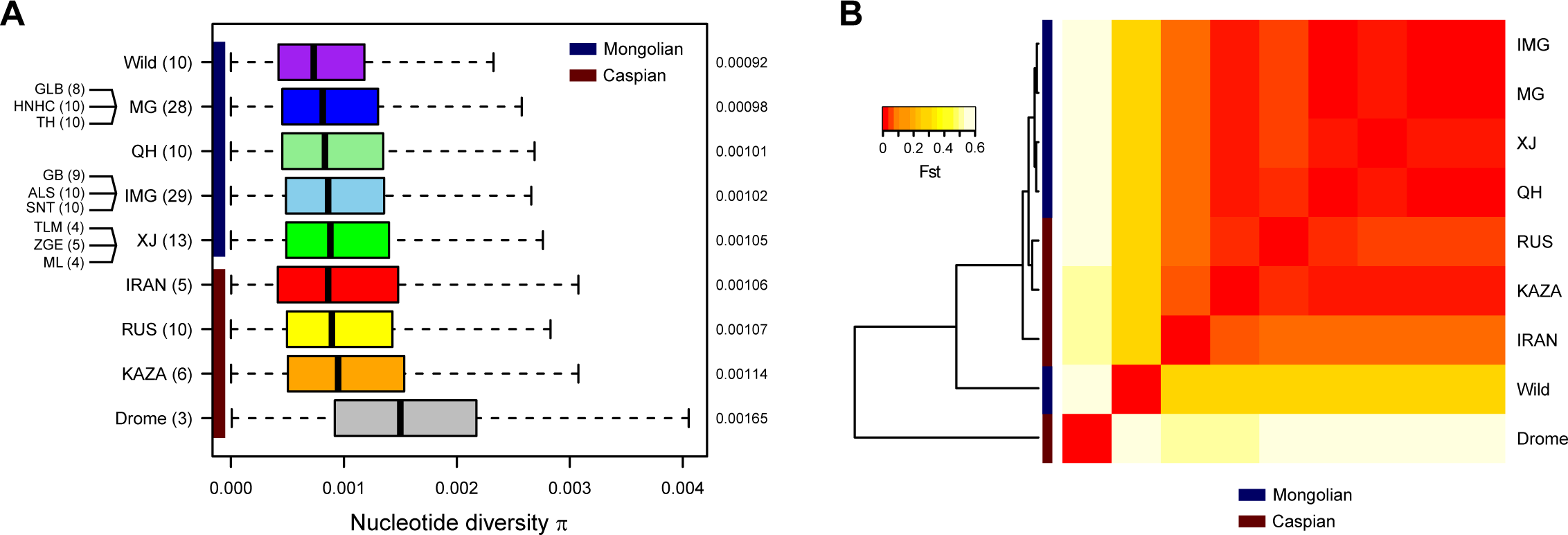
Genetic diversity and differentiation of the camel populations. (A) Nucleotide diversity π. The boxplot shows π for 10-kb sliding windows across the genome. The geographic origin and sample size of each population are shown on the left, and the average of π are shown on the right. Multiple local breeds were sampled for MG, IMG and XJ. Individuals with close genetic relationships were removed. (B) Pairwise population differentiation Fst. The heatmap represents average Fst for 10-kb sliding windows. The dendrogram represents hierarchical clustering of the populations based on Fst.

We then measured the pairwise genetic distance between the camel populations by Weir’s Fst (Figure 1B). The result was well in agreement with the known phylogeny, which indicated that the dromedaries had the highest Fst with the Bactrian camels (0.53-0.63), and the wild Bactrian camels had the second highest Fst with the domestic ones (0.27-0.30). The differentiation among the domestic Bactrian camels was much lower, implying a single-origin of them. Interestingly, among the domestic Bactrian camels, those from Iran displayed the largest divergence with all others (0.05-0.06). To validate the population differentiation, we constructed a neighbor-joining (NJ) tree for all individuals based on their pairwise identity-by-state (IBS) matrix (Supplementary Figure S6). The NJ tree also supported IRAN as the first population separated from all the other domestic Bactrian camels, implying the possibility of Iranian origin of domestic Bactrian camels.

### Population structure with admixture

To reveal the overall population structure with potential admixture, we pruned the SNPs by removing those with high linkage disequilibrium, extremely low minor allele frequency and potential functional effects. The multidimensional scaling (MDS) analysis based on the pruned subset reproduced the similar result as the full set (Figure 2A and Supplementary Figure S7). As expected, the dromedaries and wild Bactrian camels could be separated by the first and second coordinates, respectively. When the third coordinate was incorporated in the MDS, IRAN was separated from all other domestic Bactrian camels (Figure 2A).

**Figure 2.**
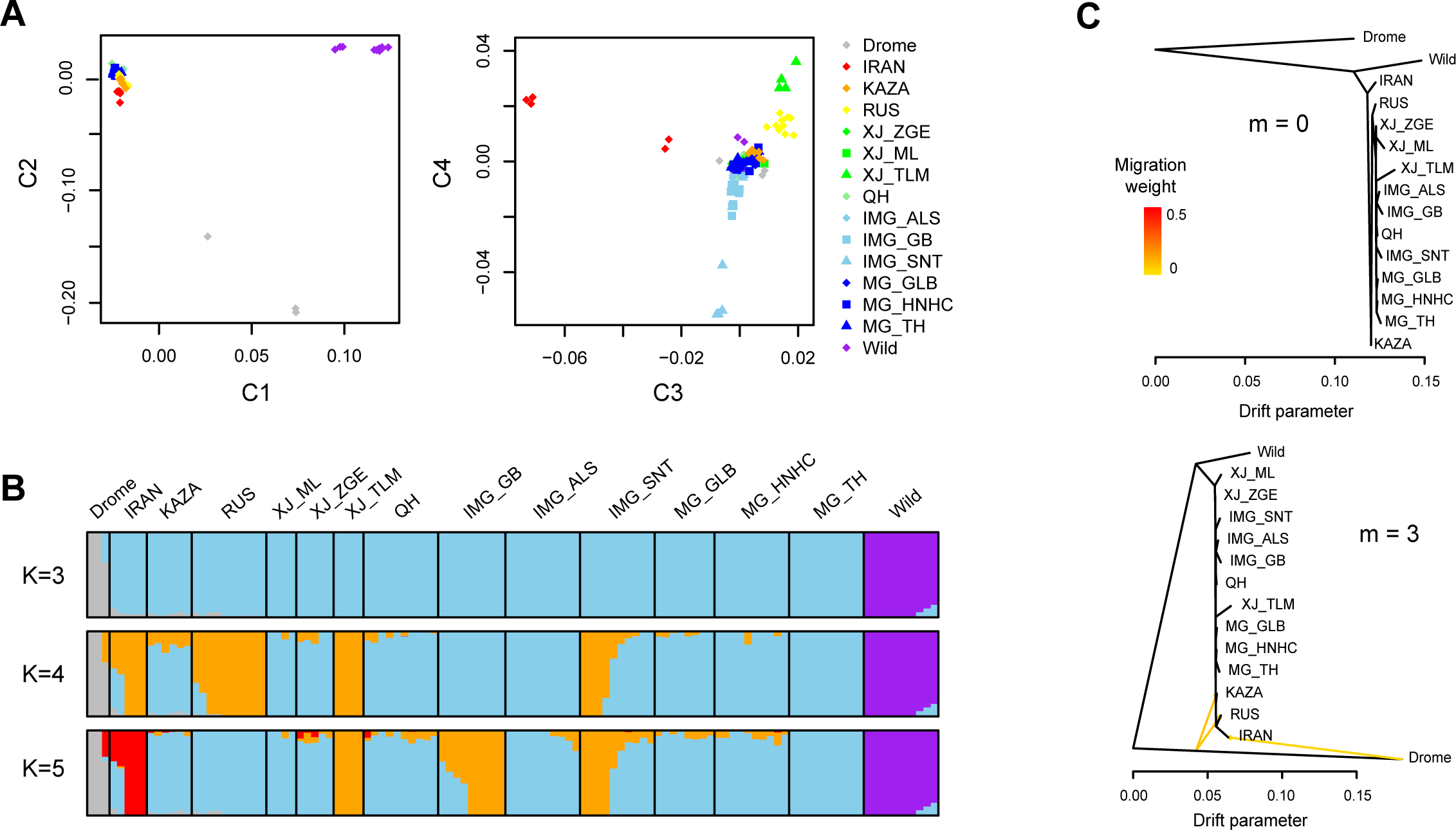
Population structure of the camels based on genome-wide SNPs. (A) Multidimensional scaling (MDS) plot with coordinate 1-4 (C1-C4). (B) Admixture analysis assuming different number of ancestry K. The proportion of an individual’s genome assigned to each ancestry is represented by different colors. (C) TreeMix analysis with different assumption of migration events m. The migration weight is the proportion of ancestry received from the donor population.

To estimate different ancestral proportions, we performed population structure analysis with Admixture [30] by assuming K ancestral populations (Figure 2B). The cross-validation procedure supported that K = 3 was optimal (Supplementary Figure S8), showing a clear division between the dromedaries, wild Bactrian camels and domestic Bactrian camels. Evident introgression of domestic Bactrian camels into the Iranian dromedaries was observed, at least in one dromedary. Accordingly, the dromedary ancestry was prevalent in the Caspian Bactrian camel populations including IRAN, KAZA and RUS, with a proportion estimated as 1%-10%. Moreover, we observed introgression of domestic Bactrian camels into several wild individuals with a proportion of 6%-15%. As the wild camels have become quite rare, it was proposed that the introgression could threaten the genetic distinctiveness of the wild species [31]. On the contrary, the wild camels contributed nearly nothing to the ancestry of domestic populations, even to the Mongolian populations which share close habitats with the wild camels. Although most domestic Bactrian camels lacked differentiation when K grew, IRAN was the first population to separate with a unique ancestry (K = 5, Figure 2B).

As another method to examine the population structure with admixture, we inferred the population tree for the camels using TreeMix [32] (Figure 2C). When no migration track (m = 0) was added, the tree topology again indicated that IRAN was the first lineage separated among all the domestic Bactrian camels. Increasing of migration tracks (m = 3) could dramatically improve the fit of the model (Supplementary Figure S9), which identified gene flows from dromedaries to domestic Bactrian camels in Middle Asia, including KAZA, RUS and IRAN with migration weights ranging from 4% to 8% (Supplementary Table S8). It was worth mentioning that while the migration track pointed at the end of the dromedary branch to IRAN, it pointed at the middle of the dromedary branch to KAZA and RUS (Figure 2C). This could imply a ghost population related to the Iranian dromedary that contributed to the ancestry of KAZA and RUS. However, TreeMix detected no signal of migration between the wild and domestic Bactrian camels. We then used the less-parameterized three- and four-population (F3/F4) test [33] to evaluate the statistical significance of the admixture events. Again, the F3 test strongly supported the admixture of dromedaries and Bactrian camels in KAZA, RUS and IRAN (Supplementary Table S9), but it did not support the admixture between the wild and domestic Bactrian camels. The more sensitive F4 test confirmed a significantly higher extent of admixture between dromedaries and Bactrian camels in Middle Asia compared with those in Northeast Asia (Supplementary Table S10). Among the latter, a higher extent of admixture with dromedaries was detected in XJ than in MG/IMG.

### Evidence for Iranian origin by removing introgression

Mongolia and Iran were the two alternative regions of domestication for Bactrian camels based on archaeological evidences [1, 12, 15], but the most probable one remained unsolved. Although we observed the largest genetic differentiation between the Iranian population and all the other domestic ones, the existence of admixture between dromedaries and Bactrian camels in Middle Asian would weaken the support for origin inference. To reduce this confounder effect, we attempted to remove the introgressed segments of dromedaries from the Bactrian camel genomes by using the ‘BABA/ABBA’ test [34]. We grouped the Mongolian and Caspian domestic populations, and compared allele sharing between the two groups with dromedaries (Figure 3A). We used the statistic f_d_, a robust version of the Patterson’s D to locate introgressed segments [35], and applied a strict significance level of Z-score = 2 by using the Jackknife procedure (Supplementary Figure S10). In a total of 21,188 non-overlapping 100-kb segments across the genome, there were far more segments with significant signals of introgression in the Caspian populations (11,651, Z > 2) than in the Mongolian populations (3,778, Z < −2) as expected. We then performed the Admixture analysis based on the remaining segments and confirmed that the introgression of dromedaries were effectively removed (Supplementary Figure S11). Re-calculation of the pairwise Fst after removing introgression still indicated that IRAN was the most differentiated one (0.04-0.06) among all the domestic populations (Figure 3B). To gain more insights into the population phylogeny, we re-constructed the NJ tree based on the pairwise Fst and performed the bootstrap test (Figure 3B). It was shown that IRAN was the first one to separate among all the domestic Bactrian populations, followed by KAZA and RUS. These results implied the Iranian origin of domestic Bactrian camels and a dispersal route from Middle Asia to Mongolia.

**Figure 3.**
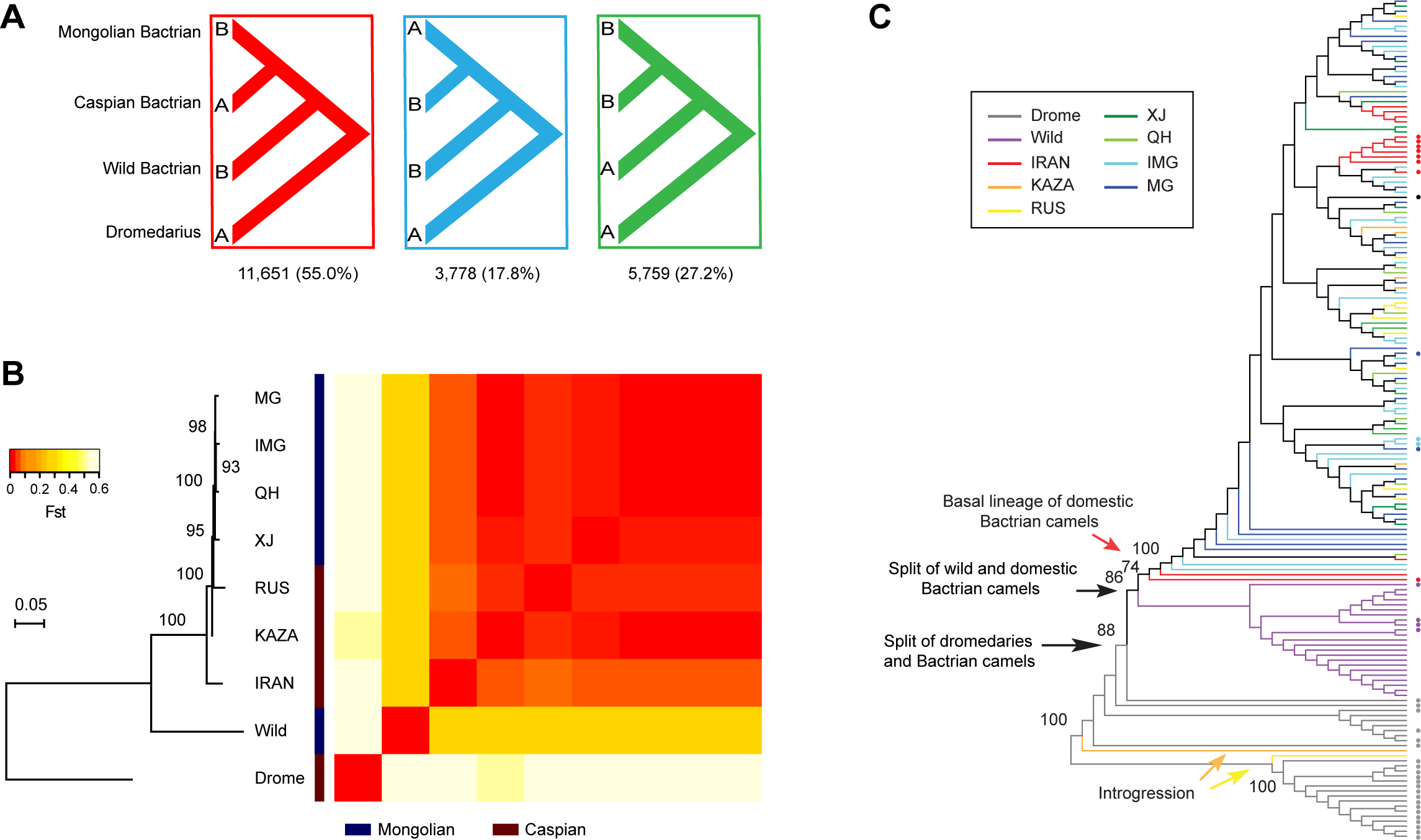
Identification of the origin of domestic Bactrian camels by removing introgression. (A) BABA/ABBA analysis for introgression of dromedaries into domestic Bactrian camels. Number of 100-kb segments with significant f_d_ for each tree configuration is shown. (B) Neighbor-joining (NJ) tree of the populations after the introgressed segments were removed. The heatmap represents average pairwise Fst for 10-kb sliding windows. Bootstrap values of the NJ tree were calculated by re-sampling the 10-kb windows for 100 times. (C) Maximum likelihood tree of full-length mtDNAs. Populations are represented by different colors, and sequences from Genbank are indicated by dots. Bootstrap values for main branches are labeled.

As an independent evidence, we also reconstructed the maximum likelihood tree of full-length mtDNAs based on the 128 samples we sequenced in this study, as well as 39 additional samples available from Genbank (Figure 3C and Supplementary Table S11). Introgression of mtDNAs could easily be identified and excluded from the tree. For example, two newly sequenced mtDNAs from KAZA and RUS were clustered with dromedaries. Within the clade of domestic Bactrian camels, although most camels from different geographic regions were mixed, two mtDNAs from IRAN formed the most basal lineages of the domestic populations (Figure 3C). Moreover, after removing the introgressed mtDNAs, IRAN displayed the highest nucleotide diversity (Supplementary Table S12). These mtDNA results were in agreement with the nuclear genome and again implicated the Iranian origin of domestic Bactrian camels.

### Demographic history of Bactrian camels

We performed several parametric modeling analyses to infer the demographic dynamics of the camels in history. Consistent with previous study [10], the long-term trajectories of Bactrian camels based on the pairwise sequentially Markovian coalescent (PSMC) model [36] revealed a tremendous decrease in the effective population size of the ancestral camels earlier than one million years ago (Supplementary Figure S12). Although the long-term trajectories of the wild and domestic Bactrian camels were generally similar, they were obvious to diverge from each other as early as 0.5 million years ago. The whole-genome divergence was in agreement with previous mtDNA studies [9, 13], which proposed that the domestic Bactrian camels and the extant wild ones were separated so long that the former could not be directly derived from the latter.

In order to explore the divergence time among the camel populations in more detail, we used the generalized phylogenetic coalescent sampler (G-PhoCS) [37]. Given the phylogeny of the camel populations, G-PhosCS could estimate the mutation-scaled population size and population divergence time based on unlinked neutral loci in individual genomes from each population (Supplementary Figure S13 and Supplementary Table S13). The time was calibrated by assuming the Bactrian camel and dromedary divergence of 5.73 million years in the Timetree database [38], When no migration band was incorporated, convergence of all parameter estimates could easily be achieved (Supplementary Figure S14 and Supplementary Table S14). Similar to the PSMC results, the effective population size was generally decreased from ancestral to modern populations (Figure 4). The divergence time between wild and domestic Bactrian camels was estimated as 0.49 million years ago (95% confidence interval [CI]: 0.35-0.65 Mya, Figure 4), which was slightly later than that based on mtDNAs (0.7 Mya) [13]. Among the domestic populations, IRAN was separated from others about 10.2 thousand years ago (95% CI: 0.25-22.5 Kya), and then the Caspian and Mongolian populations were separated about 7.39 thousand years ago (95% CI: 0.01-18.5 Kya, Figure 4). To allow for gene flow, we also tried to introduce migration bands from dromedaries to Bactrian camel populations (Supplementary Figure S13 and Supplementary Table S13). The estimates could only converge when a migration band from Iranian dromedaries to IRAN, as well as a migration band from a ghost population to KAZA were introduced (Supplementary Figure S15 and Supplementary Table S14). The total migration rate was estimated as 1% for both migration bands, though the divergence time between Bactrian camel populations was indistinguishable with the migration model.

**Figure 4.**
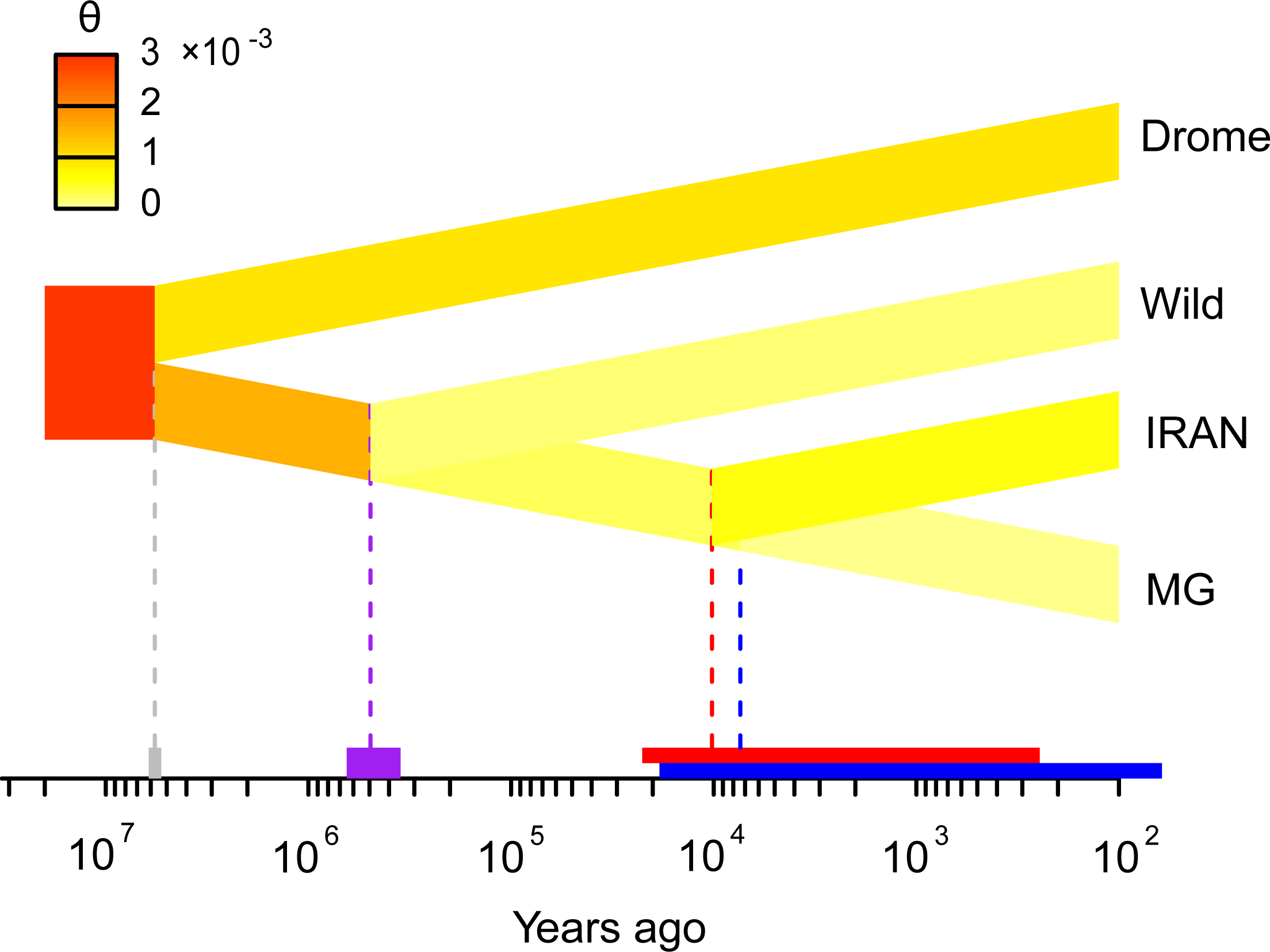
Parameter-based inference of demographic history with G-PhoCS. The change in mutation-scaled effective population size θ is represented by heat colors. The time in years were calibrated by the divergence time between dromedaries and Bactrian camels. 95% confidence intervals are shown by bars on the time axis. The red and blue bar indicates IRAN-MG and KAZA-MG divergence, respectively.

## DISCUSSION

In this study, we characterized for the first time the whole-genome variations of camels across Asia, including domestic Bactrian camels representing a major subset of recognized breeds, extant wild Bactrian camels as well as dromedaries. As the extant wild Bactrian camels are going towards extinction, our research provided extremely valuable genetic resources of the living fossil. Also, considering the extensive utilization of domestic camels in transportation, milk and wool production, our data provided new options to implement genetic association study and marker-assistant selection for improving livestock productivity and future breeding effects. In addition, these data provided an unprecedented opportunity to trace the origin and migration of domestic Bactrian camels in history. Previous studies found limited archaeological records and molecular markers for the first domestication of Bactrian camels in Middle Asia rather than in East Asia [15]. Here, we provided more evidences on the basis of the whole-genome sequences to support that hypothesis, particularly for the domestication in the area of Iran. We found that IRAN exhibited the largest genetic distance from all other domestic populations, and it harbored a unique ancestry not present in other populations. Further phylogenetic analysis revealed that IRAN was also the first one to separate among the domestic populations, which was followed by the other Caspian and Mongolian populations. Although evident introgression of dromedaries was observed in the Caspian populations, we demonstrated that it will not influence our results by removing the introgressed genomic segments. In contrast, although the extant wild and domestic Bactrian camels share close habitat in Mongolia, our whole-genome analyses gave a coherent result to other mtDNA analyses that the two populations were separated by so long a time that the latter were not likely to originate from the former [9, 13]. Furthermore, the extant wild camels contributed little to the gene pool of domestic populations, implying that the domestic populations in Mongolia could possibly be immigrated there during more recent periods.

Based on these results, we proposed a comprehensive scenario for the origin and migration of the Bactrian camels (Supplementary Figure S16). After the ancestor of camels moved from North America and split into dromedaries and Bactrian camels, the wild Bactrian camels spread from East to Middle Asia about 0.5 million years ago. The Bactrian camels were first domesticated in the area of Iran around 10 thousand years ago, which were then migrated back to East Asia with the increasing economic exchange and cooperation between West and East. In more recent periods, the domestic Bactrian camels around the Caspian Sea were further hybridized with domestic dromedaries out of Arabia. This scenario could well resolve the mystery why the wild and domestic Bactrian camels from the Mongolian Plateau have so large genetic distance. It was also similar to what happened in human evolution. Modern genomic evidence suggested that although many groups of ancient hominids occupied the continent in history, they were globally extinct and substituted by only a small group of people out of Africa [18].

Despite the insights gleaned from our data, it was important to note that the direct wild progenitor of domestic Bactrian camels were not found in Iran now, which may no longer exist. However, there were records suggesting that the wild Bactrian camels were more widely distributed throughout Asia in history, extending from the great bend of the Yellow River westward to central Kazakhstan [7]. In future work, sequencing of ancient genomes from camel fossils will add to the picture of their early domestication.

## METHODS

### Sample preparation

Blood samples of 105 domestic Bactrian camels were collected from villages in China (55), Mongolia (28), Kazakhstan (6), Russia (10) and Iran (6). Blood samples of 4 dromedaries were also collected from Iran. The collection were made during routine veterinary treatments with approval from the Camel Protection Association of Inner Mongolia. An endeavor was made to collect samples from unrelated individuals based on the information provided by the owners and local farmers. We collected 50 ml blood for each camel from the jugular vein after disinfection treatment, placed it in EDTA anticoagulant tubes, and then stored it at −80 °C. Skin samples from the ear (0.5 cm) of 19 wild Bactrian camels were collected from the Great Gobi-Strictly Protected Area “A” in Mongolia. The wild Bactrian camels chosen were artificially reared, and the research was reviewed and approved by the Great Gobi National Park. Proper surgical procedures were adopted in the collection. Local anesthesia (5% Procaine Hydrochloride) was applied to the ear, and the wound was disinfected with iodophor and sulfonamide powder. The samples were eluted with PBS buffered solutions, placed in cryotubes and stored at −80 °C.

### Genome sequencing

The genomic DNA was extracted from 200 μl blood samples with the QIAamp DNA Blood Mini kit (Qiagen), and from the skin samples with a standard phenol-chloroform method. The quality and integrity of DNA was controlled by OD260/280 ratio and agarose gel electrophoresis. For sequencing library preparation, the genomic DNA was sheared to fragments of 300–500 bp, which were then end-repaired, “A”-tailed, and ligated to Illumina sequencing adapters. The ligated products with sizes of 370–470 bp were selected on 2% agarose gels and then amplified by PCR. The libraries were sequenced on Illumina HiSeq platform with standard paired-end mode.

### Variant calling

We used an in-house script to perform quality control on raw sequencing reads. Low-quality reads marked by Illumina sequencers in the FASTQ files were removed. 3’-ends with base quality score < 20 were trimmed, and reads with length < 35-bp were removed after trimming. Trimmed reads were then mapped to the reference genome assembly of the Bactrian camel (ftp://ftp.ncbi.nih.gov/genomes/Camelus_ferus/CHR_Un/cfe_ref_CB1_chrUn.fa.gz) using BWA-MEM (v0.7.12) [39] for each individual with default parameters. We followed the GATK pipeline (v3.2-2) [40] for variant calling. Firstly, PCR duplicates were removed using Picard tools (v1.135), and local indel realignment were performed. Secondly, SNPs and small indels were called across all 128 individuals simultaneously. Finally, the raw variants were filtered with the following criteria: 1) variant quality score > 40; 2) average sequencing depth of individuals 1-40; 3) individuals with missing genotypes < 20. Total number of SNPs were reduced from 17.76 to 15.76 million after filtering. Functional annotation of variants were performed with ANNOVAR (v2013-06-21) [41] according to RefSeq (ftp://ftp.ncbi.nih.gov/genomes/Camelus_ferus/GFF/ref_CB1_scaffolds.gff3.gz).

### Population statistics and structure

Summary population statistics, including pairwise nucleotide diversity π, Watterson’s θ and Weir’s Fst across 10-kb sliding windows were calculated by VCFtools (v0.1.12b) [42]. Pairwise kinships between the samples were inferred by KING (v2.1.3) [43], and one of the paired individuals with close relationship was removed. The following SNPs were further removed by PLINK (v1.07) [44] for population structure analyses: 1) multiallelic sites; 2) SNPs in approximate linkage disequilibrium with each other (--indep 50 5 0.2); 3) SNPs with extremely low minor allele frequency (--maf 0.005); 4) SNPs located within exons and flanking 1-kb regions. A total of 2.65 million SNPs were preserved. Multidimensional scaling and pairwise distance matrix based on IBS were calculated using the ‘--mds-plot 4’ and ‘--distance-matrix’ option in PLINK, respectively. The distance matrix was used to construct the neighbour-joining tree by Phylip (v3.69) [45]. 100 random datasets were generated with ‘--thin 0.1’ option in PLINK, and bootstrap values were retrieved from the consensus tree reconstructed by Phylip. The population ancestry was inferred by ADMIXTURE (v1.3.0) [30] with a fast maximum likelihood method. The optimum number of ancestral clusters K was estimated with the 5-fold cross-validation procedure.

### TreeMix analysis and admixture tests

Migration events among camel populations were inferred using TreeMix (v1.12) [32] with migration number m = 0-5. The threepop/fourpop module from the TreeMix package was used to perform the F3/F4 test [33, 46] with ‘-k 500’. In the F3 test (Z; X, Y), one focal population (Z) was tested as a mixture of population X and Y. A large negative value of F3 score (standardized to Z-score with the Jackknife procedure) would indicate a very strong signal of Z as a mixture of X and Y. In our analysis, we ran F3 tests with all configurations of the populations. In the more sensitive F4 test (Y, Z; W, X) where W is an outgroup of Y and Z, the admixture bias between Y and Z with X was tested. If Y (or Z) have more admixture with X, it will show significant negative (or positive) F4 score (standardized to Z-score with the Jackknife procedure). To focus on the admixture between the domestic Bactrian camels and dromedaries, we set the population configuration as (Y, Z; wild, drome), where Y and Z were two domestic populations.

### Local introgression test

To select the local genomic regions with significant introgression between dromedaries and Bactrian camels after their divergence, we used an in-house script to perform the BABA/ABBA test [34] across 100-kb sliding windows. For the tree configuration (Y, Z; W, X), the original Patterson’s D statistic can be calculated as a normalized F4 score [46]:

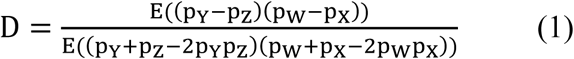

where pX is the frequency of a given allele in population X, and the expectations E() are estimated by averaging all SNPs in a window. The more robust f_d_ statistic for local genomic regions, which is a special form of F4-ratio and directly measures the proportion of introgression [35], can be formulated as:

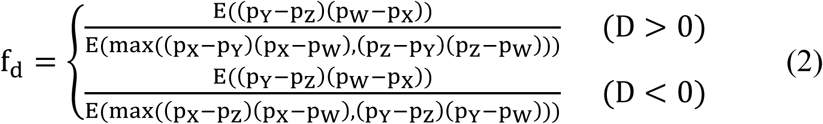

We used the population configuration (Mongolian, Caspian; wild, drome) to performed the test. The f_d_ statistic in each window was evaluated by the Z-score with the Jackknife procedure:

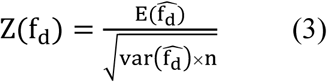

where 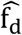 is estimated with a 10-kb block removed each time, and n is the repetition times.

### Mitochondrial DNA analysis

We complied the full-length mtDNA sequences of camels from both we sequenced in the study and those collected from GenBank. The sequences were aligned using ClustalW [47]. The control regions were deleted because they were missing in many sequences and not well aligned. The maximum likelihood tree was constructed using MEGA (v6.06) [48] with 1000 random bootstrap runs. The Tamura-Nei model and uniform substitution rates among sites were adopted. The nucleotide diversity of mtDNAs were calculated by DNASP (v5.1.0.1) [49].

### G-PhoCS analysis

To prepare for the G-PhoCS (v1.3) [37] input, we implemented the following filters to the genome to reduce the effects of selection and sequencing errors: 1) exons and 1-kb flanking regions; 2) gap regions in the genome assembly; 3) regions with repeat sequence annotations. Altogether, 47% of the genome were excluded. We then randomly collected 10,000 1-kb loci located at least 30 kb apart to ensure sufficient inter-locus recombination. Multiple sequence alignments for the loci from individual genomes per population were retrieved by vcf-consensus in VCFtools [42], with heterozygous genotypes represented by the IUPAC code and missing genotypes masked by ‘N’. Recommended gamma priors were used in the G-PhoCS analysis for the mutation-scaled population size θ, population divergence time τ, as well as migration rate m. The MCMC was run for 100,000 burn-in iterations and 500,000 sampling iterations with 10 iterations between two traced samples. The automatic fine-tuning procedure was done during the first 10,000 iterations. The convergence and mixing of the MCMC trace were monitored by Tracer (v1.6, available from http://tree.bio.ed.ac.uk/software/tracer/). Because the stochastic nature of the MCMC algorithm, we tested the models on independent datasets, and accepted the results only if two independent runs achieved the similar estimates. Simple model without migration had a high chance to converge, but none of the runs with migration bands directly from dromedaries to IRAN and/or KAZA showed convergence. This could be improved by incorporating a ghost population closely related to the dromedary and with migration band to KAZA. All loci of the ghost population were set as ‘N’. The time scale in years was calibrated according to a consensus divergence time of Bactrian camels and dromedaries, which was 5.73 million years ago in TimeTree [38]. The total migration rate across a given migration band was calculated with M = mτ_m_, where τ_m_ was the time span of the migration band.

### Data availability

The data generated from this study have been submitted to the NCBI Sequence Read Archive (http://www.ncbi.nlm.nih.gov/sra/) under accession number SRP107089. The data are also available from NODE (http://www.biosino.org/node/) under accession number OEP000024.

## ACKNOWLEDGMENTS

This work was supported by grants from the International S&T Cooperation Program of China (2015DFR30680, ky201401002), the National Natural Science Foundation of China (31360397, 31560710), the National Key R&D Program of China (2017YFA0505500, 2016YFC0901704), the special project of the Inner Mongolia Autonomous Region, and the Youth Innovation Promotion Association CAS (2017325). We thank Dr. Feng Qiu and Mr. Ze Xu from BasePair Biotechnology Co., Ltd. for technical assistance on NGS.

## AUTHOR CONTRIBUTIONS

Y.L., Z.W. and J. designed the study. S.H., G.C., T.J., N.H.E., M.B., G.K.B., T.G.E., B.T, W.Z., A.Z., H., E., A.N., P.M., N., G.M., C.N., O.K., S., S. and A. collected samples. L.M., L.Y., G.D., J.H. and L.H. performed sequencing experiments. L.M., L.Y.Y., L.Y., W.L. and Z.W. analyzed the data. L.M., L.Y.Y., Z.W. and J. wrote the manuscript.

