## supplementary for "Whole-genome sequencing of 128 camels across Asia provides insights into origin and migration of domestic Bactrian camels"

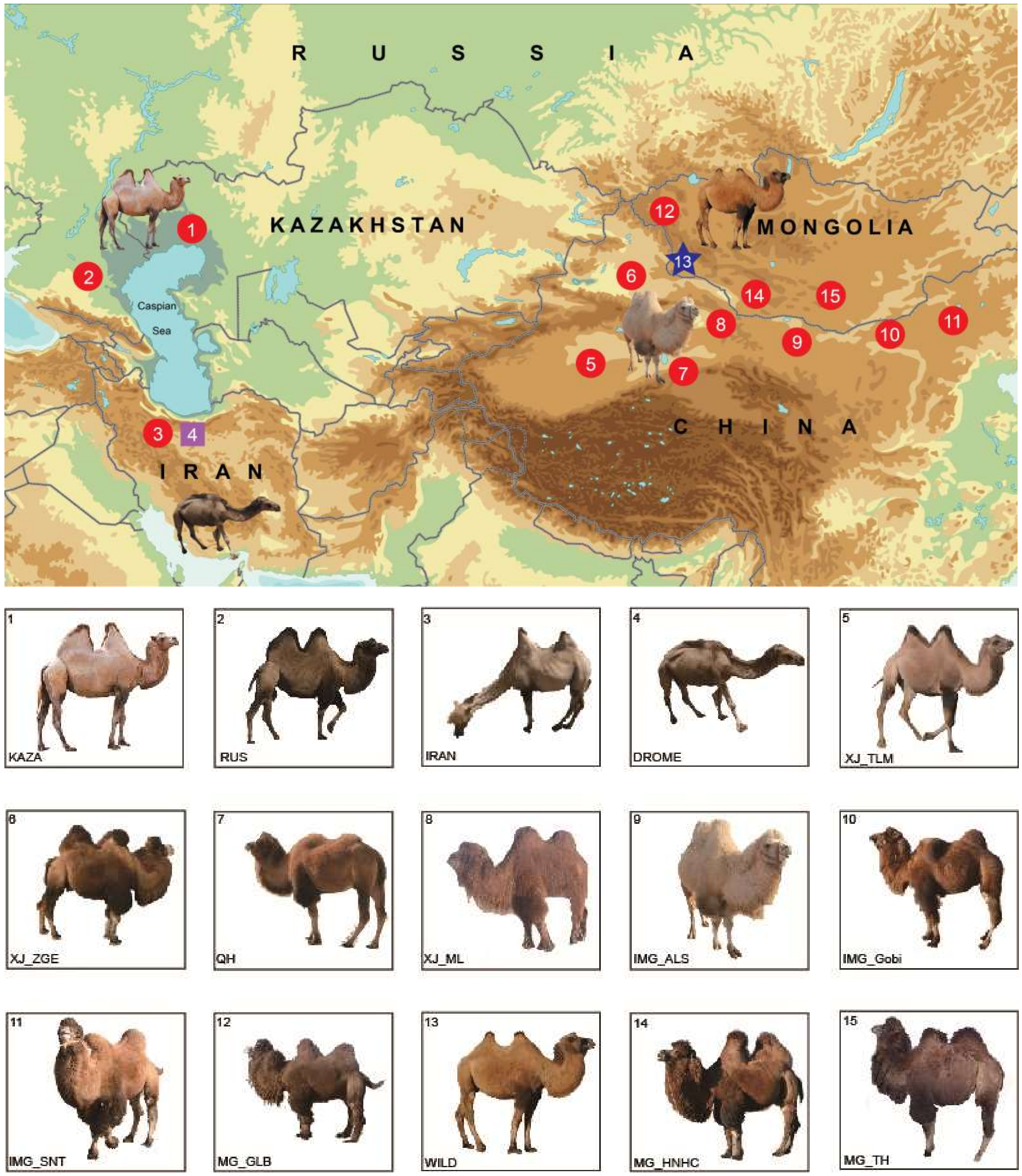


**Supplementary Figure S1. Geographic locations of the camel samples.** The Bactrian camels are mainly distributed in the Mongolian Plateau (Northeast Asia) and around the Caspian Sea (Middle Asia). Populations sampled at each location and their morphological features were shown in the bottom. The population abbreviations and sample size were listed in Supplementary Table S1.

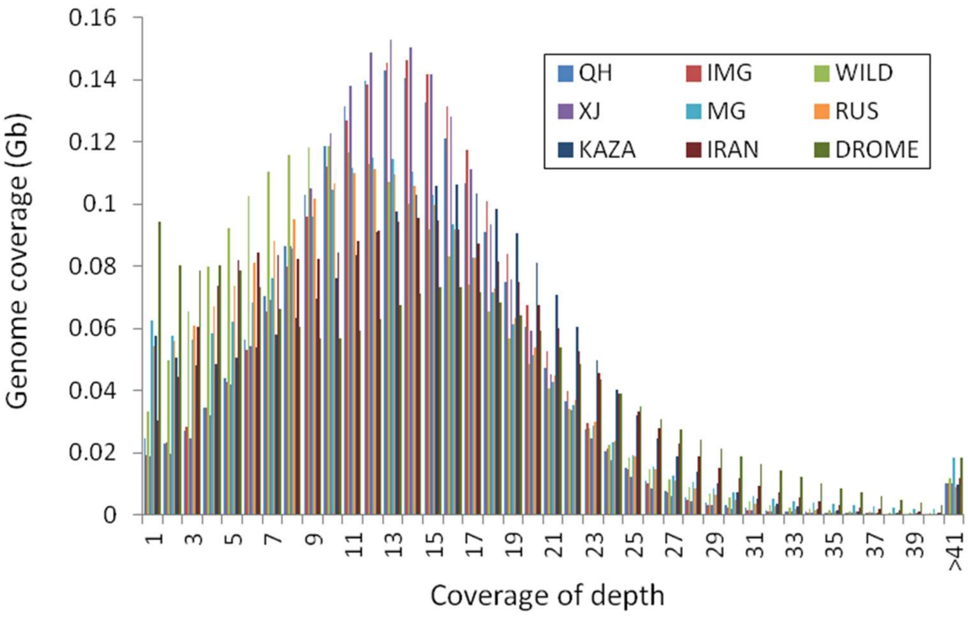


**Supplementary Figure S2. Sequencing depth distributions of the camels.** The sequencing depths of individuals from the same region were averaged for plotting. The mode of each distribution is 13× and most of the bases are covered with a depth ranging from 1 to 40×.

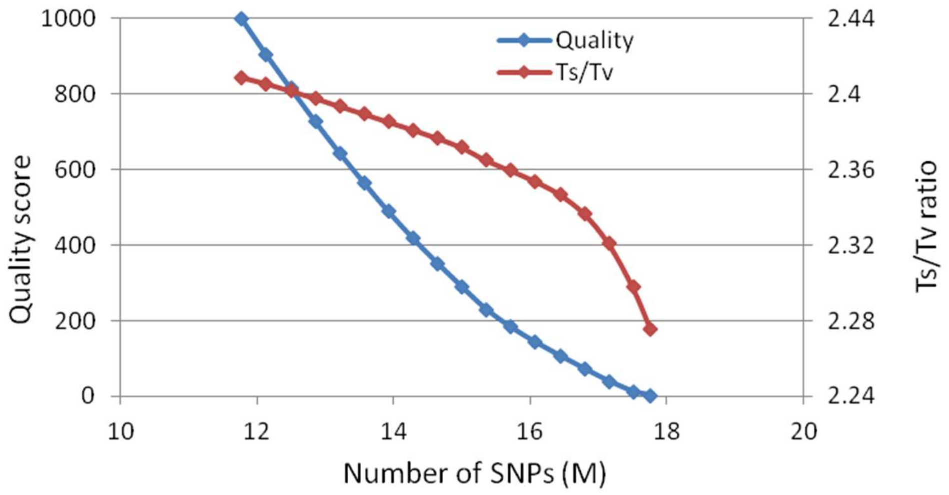


**Supplementary Figure S3. Quality of raw SNPs.** The quality score is defined as a phred score: $-10\log_{10}$Prob (variant call is wrong) in the .vcf file. Ts/Tv is the transition to transversion ratio.

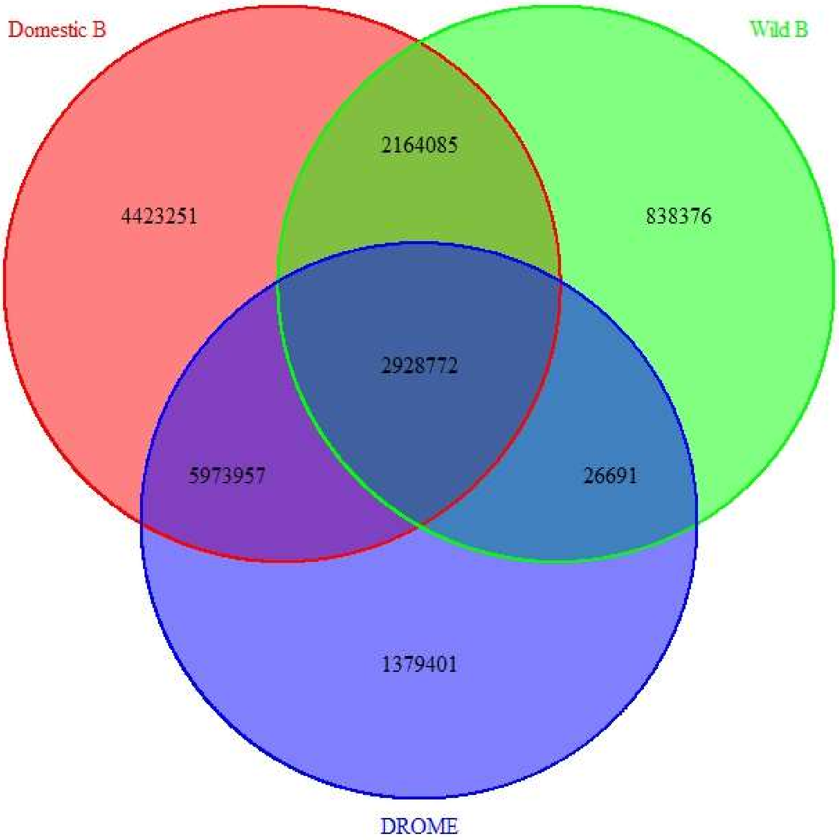


**Supplementary Figure S4. Overlapping of genomic variants between domestic Bactrian camels, wild Bactrian camels and dromedaries.**

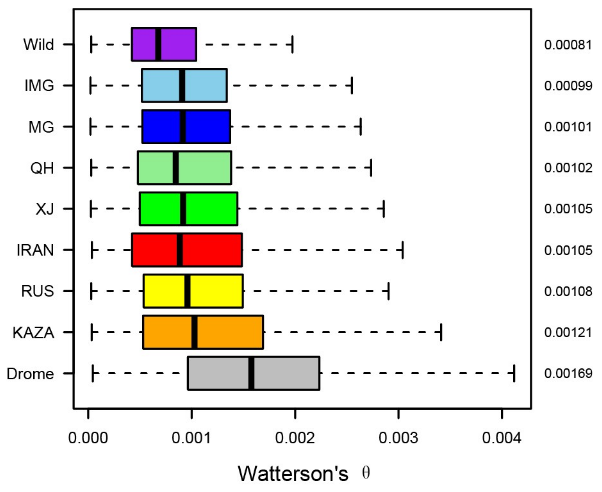


**Supplementary Figure S5. Watterson's θ of the camel populations.** The statistics were calculated with 10-kb windows across the genome. The means are shown on the right.

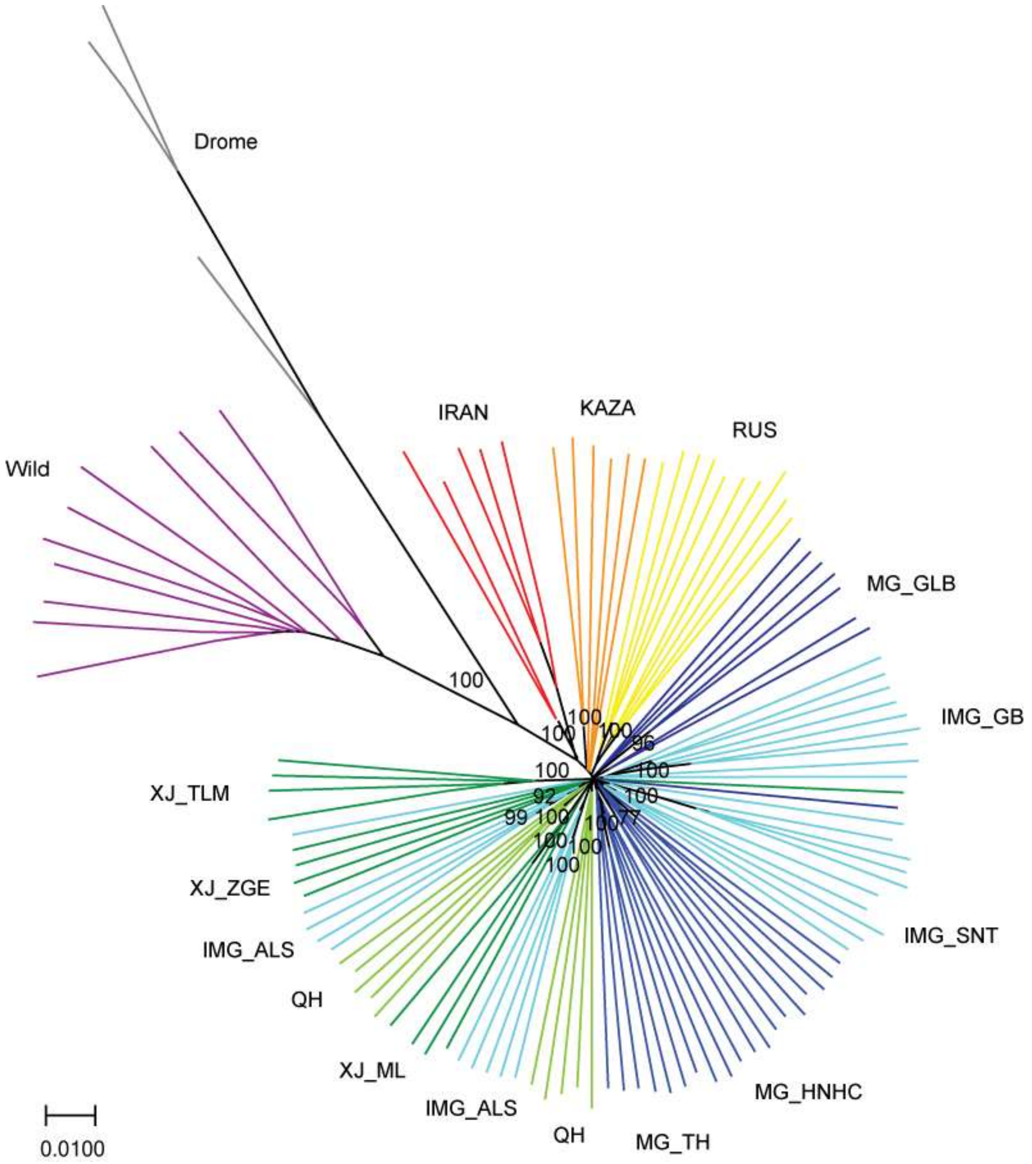


**Supplementary Figure S6. The neighbor-joining tree of all camels based on pairwise IBS.** Numbers indicate bootstrap values for main branches of the populations.

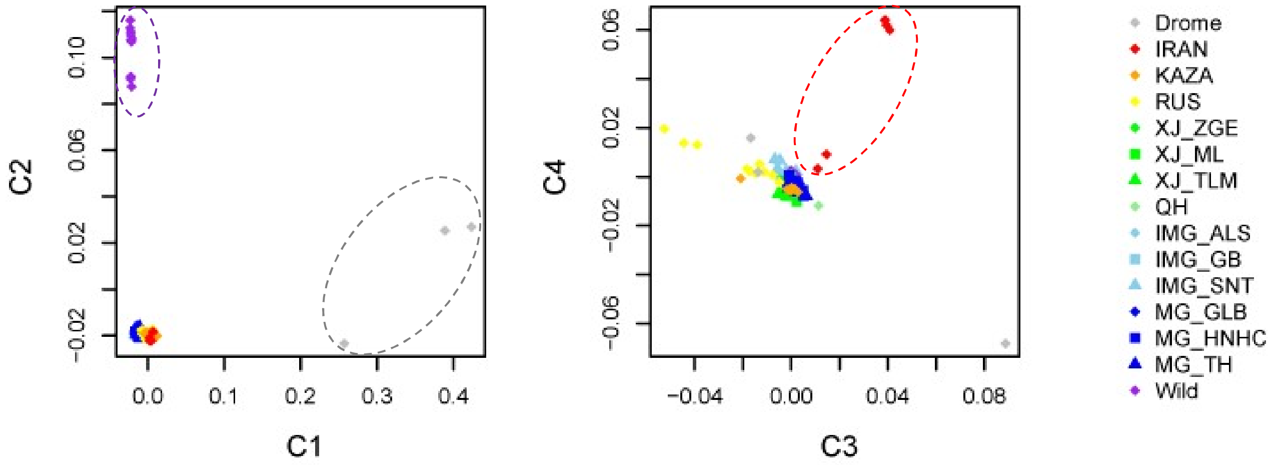


**Supplementary Figure S7. Multidimensional scaling (MDS) plot based on all SNPs.** The result is similar to that based on the pruned subset of SNPs.

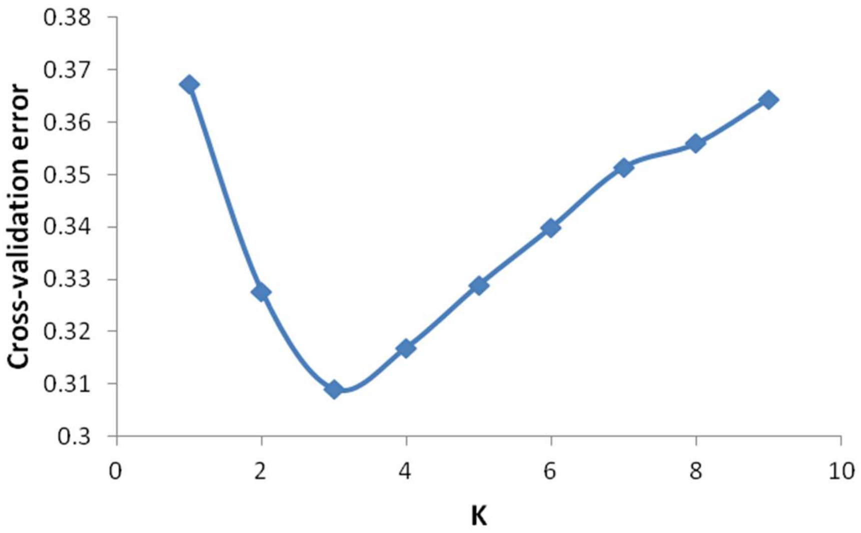


**Supplementary Figure S8. Cross-validation errors in the ADMIXTURE analysis.** The number of ancestry K was assumed from 1 to 9 and K = 3 is the optimum number.

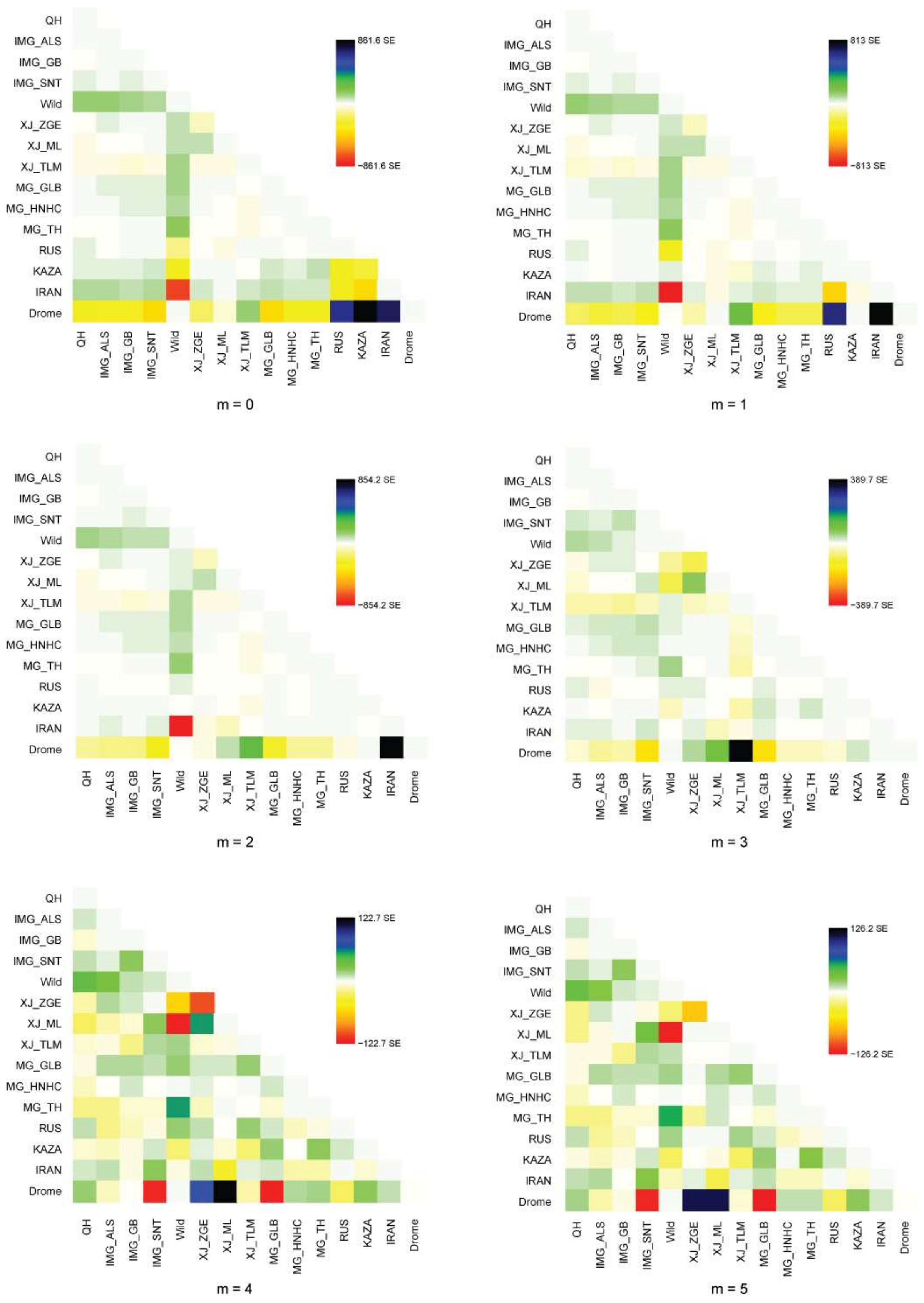


**Supplementary Figure S9. Residue visualization for the fit of TreeMix to the data with different number of migration events m**. Large positive residues indicate pairs of populations where the fit might be improved by adding additional migration edges.

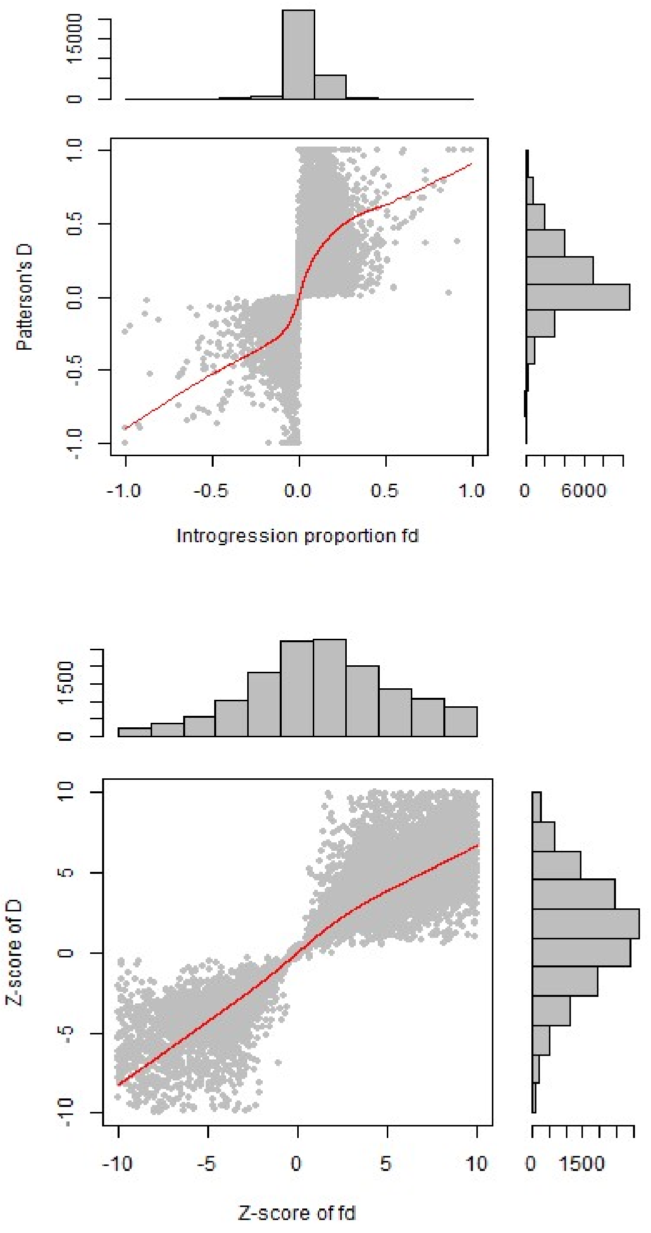


**Supplementary Figure S10. Statistics of the BABA/ABBA test.** The test was performed for 100-kb sliding windows based on population configuration (Mongolian, Caspian; wild, drome). (A) Comparison of Patterson's D and introgression proportion $f_d$. The latter shows smaller variance than the former. (B) Z-score distribution of the two statistics, which was calculated by the Jackknife procedure with a 10-kb block removed each time.

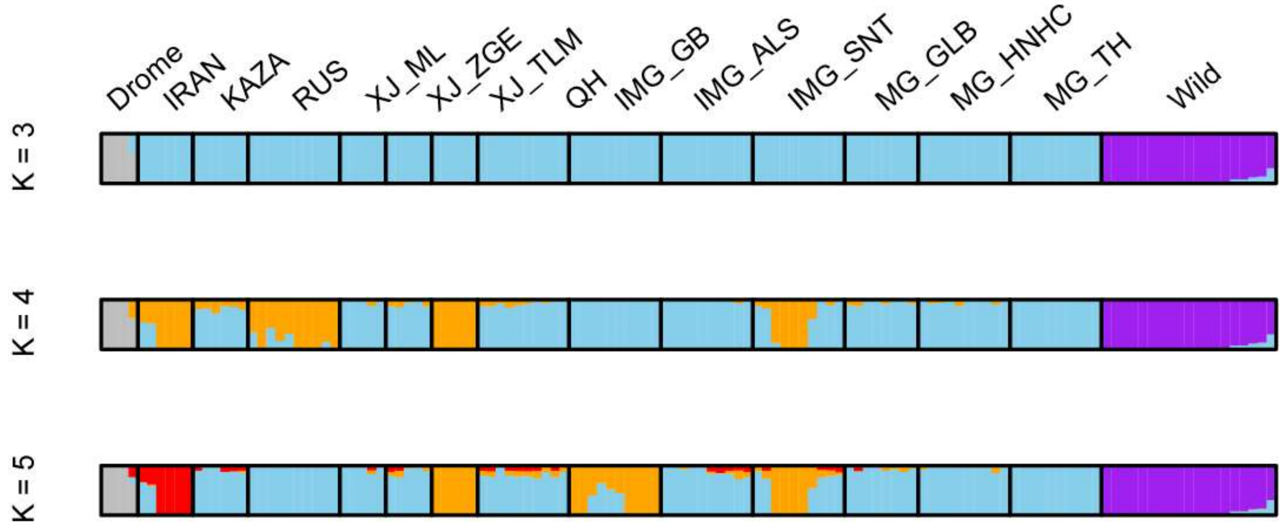


**Supplementary Figure S11. Admixture analysis after removing introgression.** The ancestry of dromedaries in IRAN/KAZA/RUS was removed by excluding genomic segments showing Z-score > 2 with the local BABA/ABBA test.

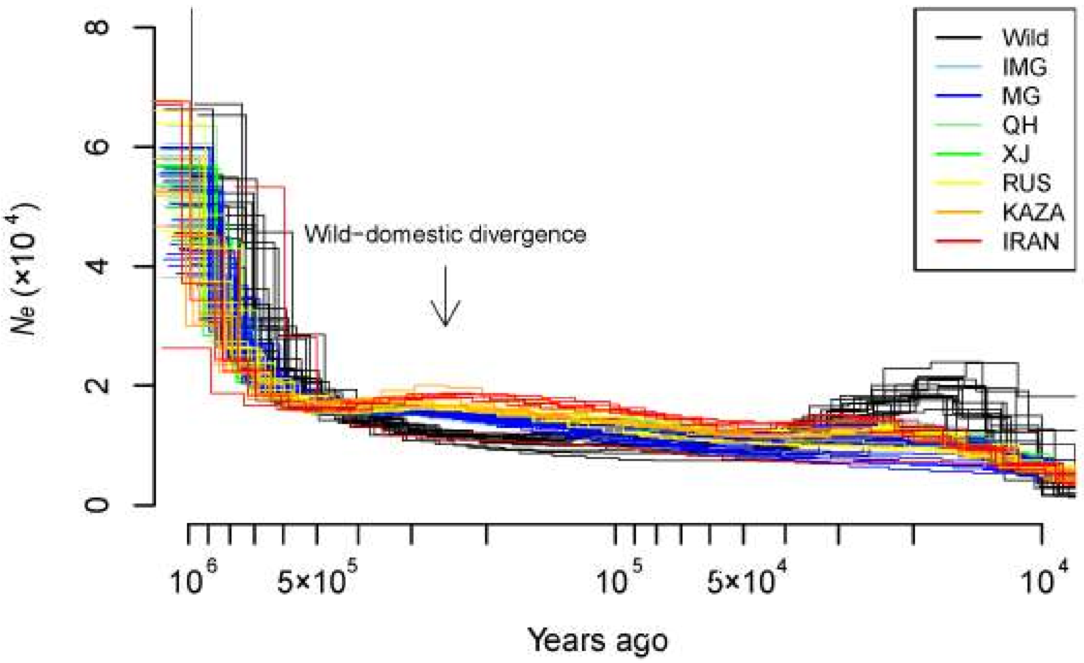


**Supplementary Figure S12. Historical effective population size (*N*e) of Bactrian camels inferred by PSMC.** Each trajectory is based on one individual genome. The arrow indicates the separation of trajectories between wild and domestic Bactrian camels. To comply with previous studies, the results were scaled with a generation time of 5 years and a mutation rate of $2.5\times10^{-8}$ per site per generation.

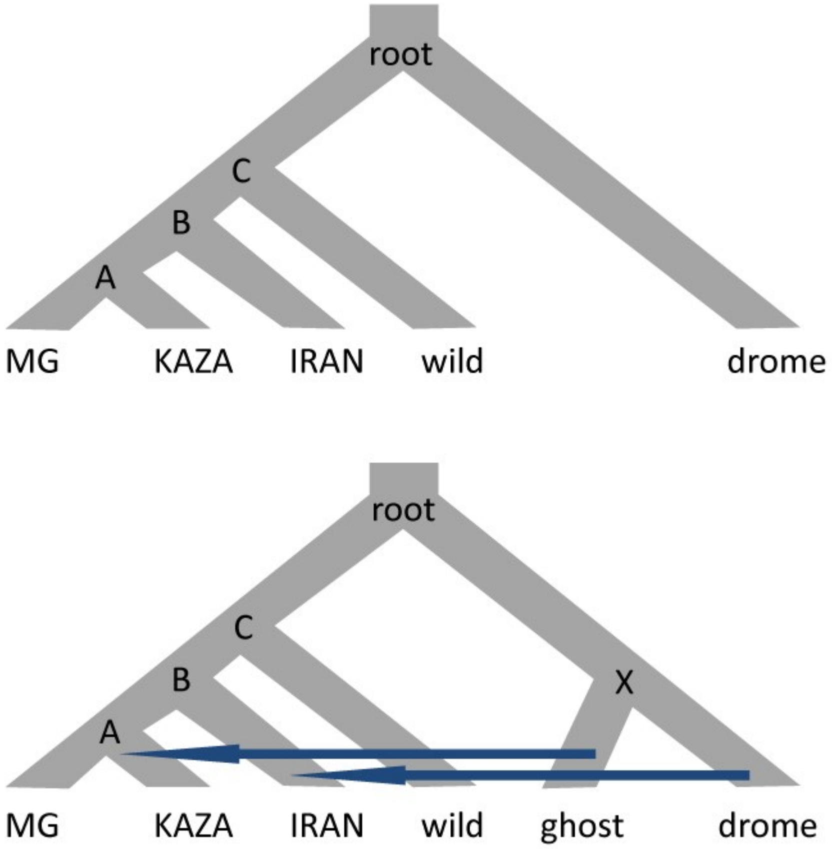


| Population | Sample |
|---|---|
| MG | SAMN06759179 |
| KAZA | SAMN06759203 |
| IRAN | SAMN06759205 |
| Wild | SAMN06759139 |
| Drome | SAMN06759213 |

**Supplementary Figure S13. Population phylogeny used in the G-PhoCS analysis.** (A) Phylogeny without migration based on the Fst distance. Ancestral populations were labeled in the tree. (B) Migration bands from the dromedary to IRAN, and a ghost population closely related to the dromedary to KAZA. (C) One diploid sample per population was used for G-PhoCS analysis.

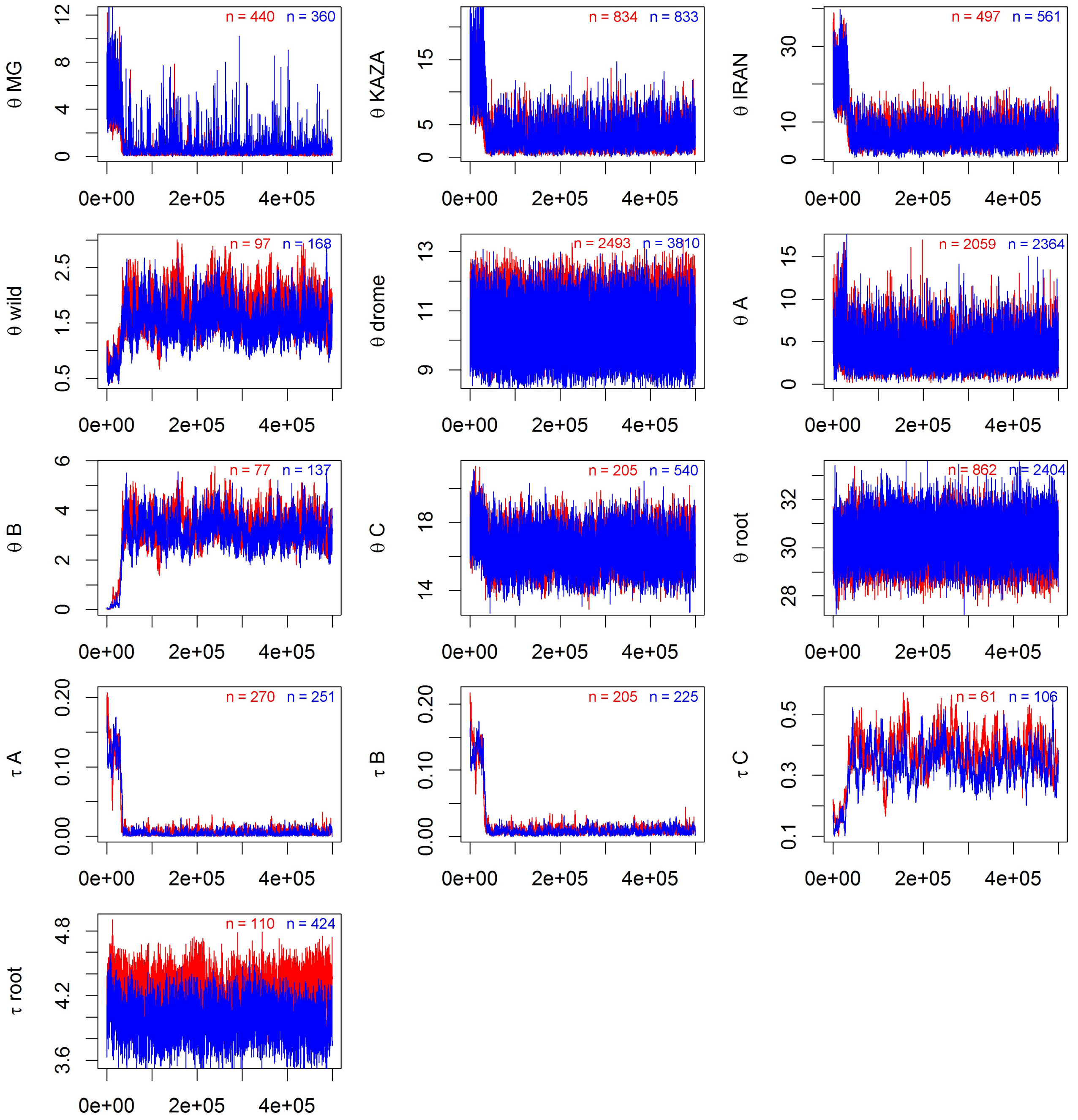


**Supplementary Figure S14. MCMC traces in the G-PhoCS analysis without migration.** θ and τ are the mutation-scaled effective population size and population divergence time, respectively (scaled up by $10^4$). A total of 500,000 iterations with 10 iterations between two traced samples are shown. Two independent runs on independent datasets are represented by two different colors. n is the effective sample size with auto-correlation adjustment by Tracer. The traces were checked for convergence and mixing of MCMC, and the first 1/10 traced samples were discarded.

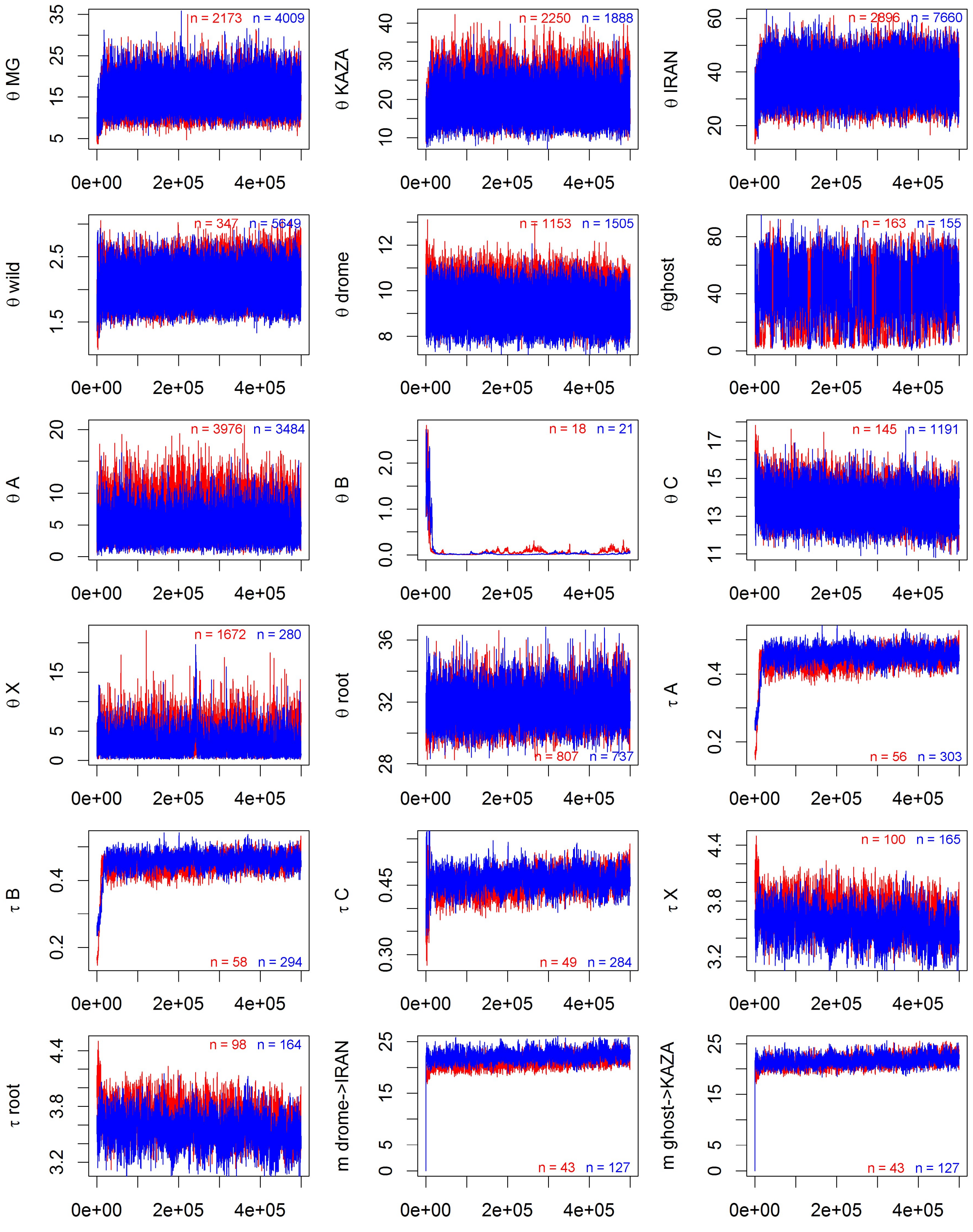


**Supplementary Figure S15. MCMC traces in the G-PhoCS analysis with migration.** m is the mutation-scaled migration rate per generation (scaled by 0.1). The first 1/10 traced samples were discarded.

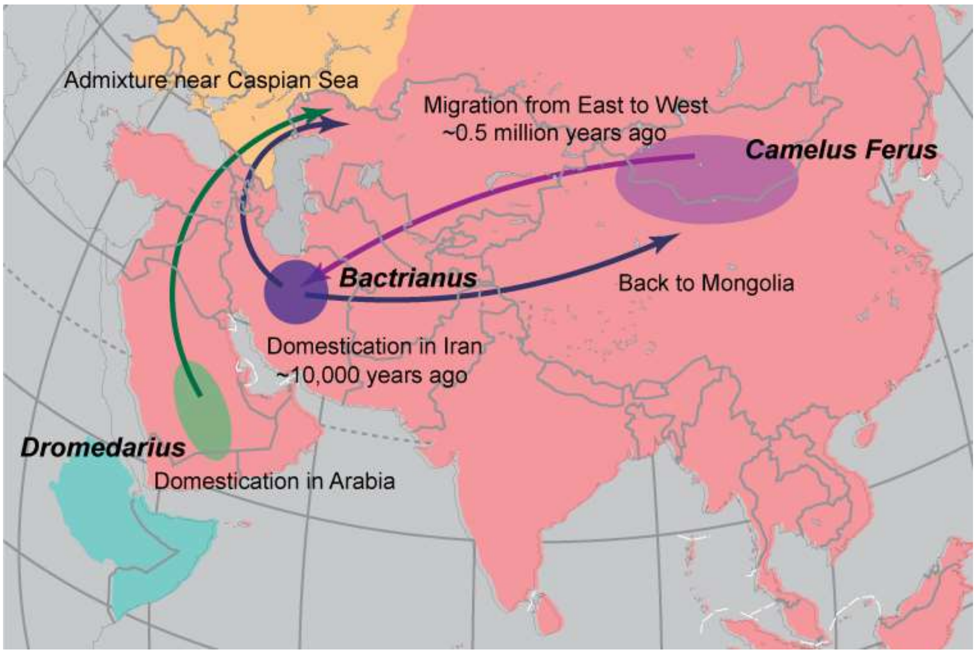


**Supplementary Figure S16. A proposed migration route of Bactrian camels across Asia based on the results from our study.**

**Supplementary Table S1.Sample information and summary of sequencing data.**

| Country | District | Breed (Abbr.) | NCBI sample accession (PRJNA383081) | Sample size | Mean raw bases (Gb) | Mean bases after QC (Gb) |
|---|---|---|---|---|---|---|
| China | Qinghai Province | Qinghai (QH) | SAMN06759087-06759096 | 10 | 32.27 | 32.10 |
| China | Alashan, Inner Mongolia | Alashan (IMG_ALS) | SAMN06759097-06759106 | 10 | 33.34 | 33.24 |
| China | Bayan Nur, Inner Mongolia | Gobi Red (IMG_GB) | SAMN06759107-06759116 | 10 | 33.16 | 33.09 |
| China | Xilingol, Inner Mongolia | Sonid (IMG_SNT) | SAMN06759117-06759126 | 10 | 32.85 | 32.79 |
| Mongolia | Gobi Altai | Wild | SAMN06759127-06759145 | 19 | 37.24 | 33.22 |
| China | Altay Fuyun, XinJiang | Zhungeer (XJ_ZGE) | SAMN06759146-06759150 | 5 | 34.91 | 34.55 |
| China | Mulei Kazakh, XinJiang | Mulei (XJ_ML) | SAMN06759151-06759155 | 5 | 31.51 | 31.43 |
| China | Tarim Basin, XinJiang | Tarim (XJ_TLM) | SAMN06759156-06759160 | 5 | 30.60 | 30.55 |
| Mongolia | Khanbogd and Bayan-ovoo soums, Umnugobi Province | Galbiin Goviin ulaan (MG_GLB) | SAMN06759161-06759168 | 8 | 30.99 | 30.91 |
| Mongolia | Mandal-ovoo soum, Umnugobi Province | Khaniiin khetsiin khuren (MG_HNHC) | SAMN06759169-06759178 | 10 | 32.70 | 32.64 |
| Mongolia | Togrog soum, Gobi-Altai Province | Tokhom-tungalag (MG_TH) | SAMN06759179-06759188 | 10 | 31.44 | 31.38 |
| Russia | The Republic of Tyva, Kalmykia | Kalmyk (RUS) | SAMN06759189-06759198 | 10 | 30.79 | 30.69 |
| Kazakhstan | Almaty Province | Kazakhstan (KAZA) | SAMN06759199-06759204 | 6 | 32.40 | 32.38 |
| Iran | Ardabili | Iran (IRAN) | SAMN06759205-06759210 | 6 | 39.19 | 33.54 |
| Iran | Ardabili | Dromedary | SAMN06759211-06759214 | 4 | 41.73 | 35.60 |

**Supplementary Table S2. Means of genome mapping statistics.**

| Breed | Mapped reads (M) | Mapped read pairs (M) | Unique read pairs (M) | Properly mapped bases (Gb) | Genome depth | Genome coverage |
|---|---|---|---|---|---|---|
| QH | 256.51 | 126.29 | 120.52 | 27.54 | 13.70 | 0.96 |
| IMG_ALS | 265.42 | 130.70 | 124.40 | 28.51 | 14.19 | 0.98 |
| IMG_GB | 264.07 | 130.16 | 123.36 | 28.30 | 14.09 | 0.98 |
| IMG_SNT | 261.68 | 128.97 | 122.94 | 28.32 | 14.09 | 0.97 |
| Wild | 337.48 | 161.20 | 150.09 | 25.96 | 12.92 | 0.98 |
| XJ_ZGE | 278.13 | 137.07 | 131.26 | 30.27 | 15.07 | 0.98 |
| XJ_ML | 250.90 | 123.52 | 117.56 | 26.74 | 13.31 | 0.96 |
| XJ_TLM | 243.90 | 120.08 | 114.99 | 26.17 | 13.03 | 0.97 |
| MG_GLB | 246.83 | 121.25 | 114.34 | 25.85 | 12.86 | 0.96 |
| MG_HNHC | 260.55 | 128.19 | 120.21 | 27.28 | 13.58 | 0.94 |
| MG_TH | 250.58 | 123.10 | 114.81 | 25.79 | 12.84 | 0.93 |
| RUS | 244.99 | 120.60 | 112.93 | 25.17 | 12.53 | 0.95 |
| KAZA | 258.27 | 127.40 | 122.79 | 28.37 | 14.12 | 0.94 |
| IRAN | 265.42 | 129.21 | 104.43 | 28.02 | 13.95 | 0.97 |
| Drome | 284.05 | 137.59 | 117.32 | 31.08 | 15.47 | 0.94 |

**Supplementary Table S3. Number of SNPs and indels after filtering.**

| Breed | Total variants | SNPs | Transition/Tranversion ratio | Small indels |
|---|---|---|---|---|
| QH | 7,581,384 | 6,751,721 | 2.4969 | 829,663 |
| IMG_ALS | 8,296,833 | 7,330,126 | 2.4619 | 966,707 |
| IMG_GB | 8,111,818 | 7,129,592 | 2.4455 | 982,226 |
| IMG_SNT | 8,135,925 | 7,159,158 | 2.4533 | 976,767 |
| Wild | 5,957,924 | 5,317,666 | 2.4371 | 640,258 |
| XJ_ZGE | 7,921,144 | 6,917,069 | 2.4332 | 1,004,075 |
| XJ_ML | 6,917,653 | 6,076,166 | 2.4544 | 841,487 |
| XJ_TLM | 6,803,884 | 5,947,454 | 2.4264 | 856,430 |
| MG_GLB | 7,750,374 | 6,867,371 | 2.4756 | 883,003 |
| MG_HNHC | 7,834,000 | 6,970,701 | 2.4968 | 863,299 |
| MG_TH | 7,082,241 | 6,324,828 | 2.5044 | 757,413 |
| RUS | 8,428,659 | 7,536,540 | 2.4922 | 892,119 |
| KAZA | 8,541,210 | 7,603,345 | 2.4906 | 937,865 |
| IRAN | 7,776,857 | 6,801,855 | 2.4886 | 975,002 |
| Drome | 10,308,821 | 9,236,526 | 2.5315 | 1,072,295 |
| All | 17,734,533 | 15,762,069 | 2.4307 | 1,972,464 |

**Supplementary Table S4. Proportion of annotated SNPs in different genomic regions.**

| Breed | Intergenic[a] | ncRNA[b] | UTR[c] | Intronic | Splicing | Exonic[d] | | |
|---|---|---|---|---|---|---|---|---|
| | | | | | | Synonymous | Non-synonymous | Stop altering[e] |
| QH | 69.71% | 0.12% | 0.17% | 29.11% | 0.0031% | 0.51% | 0.38% | 0.0042% |
| IMG_ALS | 64.21% | 0.14% | 0.20% | 34.49% | 0.0032% | 0.55% | 0.40% | 0.0043% |
| IMG_GB | 69.22% | 0.12% | 0.17% | 29.67% | 0.0027% | 0.47% | 0.34% | 0.0035% |
| IMG_SNT | 68.76% | 0.12% | 0.17% | 30.10% | 0.0027% | 0.48% | 0.36% | 0.0037% |
| Wild | 68.76% | 0.14% | 0.17% | 30.00% | 0.0034% | 0.52% | 0.40% | 0.0043% |
| XJ_ZGE | 66.05% | 0.13% | 0.19% | 32.72% | 0.0032% | 0.52% | 0.38% | 0.0039% |
| XJ_ML | 73.06% | 0.10% | 0.15% | 25.95% | 0.0024% | 0.42% | 0.31% | 0.0031% |
| XJ_TLM | 69.73% | 0.12% | 0.17% | 29.18% | 0.0025% | 0.45% | 0.34% | 0.0035% |
| MG_GLB | 75.63% | 0.09% | 0.14% | 23.45% | 0.0023% | 0.39% | 0.29% | 0.0031% |
| MG_HNHC | 70.21% | 0.11% | 0.17% | 28.65% | 0.0028% | 0.49% | 0.36% | 0.0038% |
| MG_TH | 70.38% | 0.12% | 0.17% | 28.45% | 0.0031% | 0.50% | 0.37% | 0.0039% |
| RUS | 65.90% | 0.13% | 0.19% | 32.80% | 0.0031% | 0.56% | 0.40% | 0.0039% |
| KAZA | 66.31% | 0.14% | 0.18% | 32.43% | 0.0032% | 0.54% | 0.39% | 0.0037% |
| IRAN | 67.41% | 0.12% | 0.18% | 31.43% | 0.0030% | 0.50% | 0.35% | 0.0035% |
| Drome | 66.46% | 0.13% | 0.19% | 32.29% | 0.0032% | 0.55% | 0.37% | 0.0034% |

[a] Including "intergenic", "upstream" and "downstream" given by ANNOVAR.

[b] Including "ncRNA_exonic", "ncRNA_intronic", "ncRNA_splicing" and "ncRNA_UTR".

[c] Including "UTR5” and "UTR3".

[d] Including "exonic" and "exonic;splicing".

[e] Including "stopgain SNV" and "stoploss SNV".

**Supplementary Table S5. Proportion of annotated small indels in different genomic regions.**

| Breed | Intergenic[a] | ncRNA[b] | UTR[c] | Intronic | Splicing | Exonic[d] | | |
|---|---|---|---|---|---|---|---|---|
| | | | | | | Frameshift[e] | Non-frameshift[f] | Stop altering[g] |
| QH | 67.44% | 0.09% | 0.26% | 32.04% | 0.0067% | 0.06% | 0.10% | 0.0021% |
| IMG_ALS | 67.88% | 0.09% | 0.26% | 31.62% | 0.0068% | 0.05% | 0.09% | 0.0016% |
| IMG_GB | 68.03% | 0.09% | 0.26% | 31.49% | 0.0063% | 0.05% | 0.08% | 0.0016% |
| IMG_SNT | 68.23% | 0.09% | 0.26% | 31.28% | 0.0058% | 0.05% | 0.09% | 0.0018% |
| Wild | 67.40% | 0.10% | 0.27% | 32.00% | 0.0071% | 0.08% | 0.14% | 0.0025% |
| XJ_ZGE | 68.73% | 0.08% | 0.25% | 30.82% | 0.0056% | 0.05% | 0.07% | 0.0018% |
| XJ_ML | 68.18% | 0.08% | 0.25% | 31.34% | 0.0075% | 0.05% | 0.08% | 0.0018% |
| XJ_TLM | 68.71% | 0.09% | 0.25% | 30.82% | 0.0051% | 0.05% | 0.08% | 0.0015% |
| MG_GLB | 67.94% | 0.09% | 0.27% | 31.55% | 0.0072% | 0.06% | 0.09% | 0.0019% |
| MG_HNHC | 67.52% | 0.09% | 0.27% | 31.94% | 0.0073% | 0.06% | 0.10% | 0.0021% |
| MG_TH | 67.34% | 0.09% | 0.29% | 32.10% | 0.0086% | 0.07% | 0.11% | 0.0026% |
| RUS | 67.58% | 0.09% | 0.29% | 32.06% | 0.0072% | 0.06% | 0.10% | 0.0023% |
| KAZA | 67.67% | 0.10% | 0.26% | 31.82% | 0.0067% | 0.06% | 0.09% | 0.0020% |
| IRAN | 67.09% | 0.09% | 0.26% | 32.43% | 0.0071% | 0.04% | 0.08% | 0.0016% |
| Drome | 65.81% | 0.10% | 0.28% | 33.65% | 0.0074% | 0.06% | 0.09% | 0.0024% |

[a] Including "intergenic", "upstream" and "downstream" given by ANNOVAR.

[b] Including "ncRNA_exonic", "ncRNA_intronic", "ncRNA_splicing" and "ncRNA_UTR".

[c] Including "UTR5” and "UTR3".

[d] Including "exonic" and "exonic;splicing".

[e] Including "frameshift deletion", "frameshift insertion" and "frameshift substitution".

[f] Including "nonframeshift deletion", "nonframeshift insertion" and "nonframeshift substitution".

[g] Including "stopgain SNV" and "stoploss SNV".

**Supplementary Table S6. Samples with close relationship inferred by KING.**

| Breed | Removed sample | Preserved sample | SNP number | Proportion of zero IBS | Kinship coefficient |
|---|---|---|---|---|---|
| Wild | SAMN06759128 | SAMN06759129 | 15,738,057 | 0.0008 | 0.2375 |
| Wild | SAMN06759128 | SAMN06759134 | 15,544,506 | 0.0033 | 0.195 |
| Wild | SAMN06759128 | SAMN06759137 | 15,241,706 | 0.0103 | 0.1817 |
| Wild | SAMN06759128 | SAMN06759144 | 15,702,771 | 0.0021 | 0.2777 |
| Wild | SAMN06759128 | SAMN06759145 | 15,458,755 | 0.0034 | 0.1961 |
| Wild | SAMN06759129 | SAMN06759130 | 15,679,356 | 0.0013 | 0.2561 |
| Wild | SAMN06759136 | SAMN06759137 | 15,249,506 | 0.0027 | 0.2514 |
| Wild | SAMN06759136 | SAMN06759143 | 15,710,144 | 0.01 | 0.1781 |
| Wild | SAMN06759136 | SAMN06759127 | 15,693,317 | 0.0022 | 0.2645 |
| Wild | SAMN06759137 | SAMN06759131 | 14,624,828 | 0.0032 | 0.2402 |
| Wild | SAMN06759140 | SAMN06759133 | 15,710,342 | 0.0018 | 0.2536 |
| Wild | SAMN06759140 | SAMN06759134 | 15,549,600 | 0.0027 | 0.2324 |
| Wild | SAMN06759140 | SAMN06759145 | 15,459,676 | 0.0027 | 0.2319 |
| Wild | SAMN06759141 | SAMN06759132 | 15,478,265 | 0.0033 | 0.2315 |
| Wild | SAMN06759142 | SAMN06759135 | 15,466,083 | 0.0064 | 0.2429 |
| Wild | SAMN06759142 | SAMN06759131 | 14,890,858 | 0.0087 | 0.1981 |
| Wild | SAMN06759143 | SAMN06759127 | 15,689,057 | 0.002 | 0.2459 |
| Wild | SAMN06759143 | SAMN06759139 | 15,706,944 | 0.0012 | 0.2569 |
| Wild | SAMN06759145 | SAMN06759134 | 15,324,008 | 0.0006 | 0.428 |
| IMG_GB | SAMN06759107 | SAMN06759108 | 15,731,573 | 0.0028 | 0.2417 |
| XJ_ML | SAMN06759153 | SAMN06759152 | 15,549,428 | 0.0089 | 0.1783 |
| XJ_TLM | SAMN06759159 | SAMN06759156 | 15,653,821 | 0.001 | 0.2584 |
| XJ_TLM | SAMN06759159 | SAMN06759158 | 15,716,807 | 0.001 | 0.2408 |
| IRAN | SAMN06759209 | SAMN06759207 | 15,662,682 | 0.0007 | 0.2323 |
| Drome | SAMN06759214 | SAMN06759212 | 14,407,046 | 0.0002 | 0.3205 |
| Drome | SAMN06759214 | SAMN06759213 | 14,093,224 | 0.0119 | 0.1775 |

**Supplementary Table S7. P-values of pairwise t-test for nucleotide diversity.**

| | KAZA | RUS | IRAN | XJ | IMG | QH | MG | Wild |
|---|---|---|---|---|---|---|---|---|
| Drome | 0 (49.75)[a] | 0 (58.66) | 0 (59.76) | 0 (60.93) | 0 (67.72) | 0 (67.84) | 0 (72.39) | 0 (77.10) |
| KAZA | - | 2.54E-13 (7.32) | 5.27E-19 (8.91) | 1.02E-21 (9.58) | 1.35E-47 (14.51) | 2.65E-57 (15.98) | 7.40E-83 (19.33) | 1.54E-142 (25.52) |
| RUS | - | - | 8.34E-2 (1.73) | 1.89E-2 (2.35) | 6.60E-13 (7.19) | 4.65E-19 (8.92) | 2.19E-34 (12.24) | 2.27E-79 (18.91) |
| IRAN | - | - | - | 5.56E-1 (0.59) | 1.22E-7 (5.29) | 1.62E-12 (7.07) | 9.03E-25 (10.28) | 3.99E-64 (16.94) |
| XJ | - | - | - | - | 2.18E-6 (4.74) | 5.86E-11 (6.55) | 1.23E-22 (9.80) | 2.66E-61 (16.55) |
| IMG | - | - | - | - | - | 3.98E-2 (2.06) | 9.61E-8 (5.33) | 3.76E-36 (12.57) |
| QH | - | - | - | - | - | - | 1.95E-3 (3.10) | 2.76E-24 (10.17) |
| MG | - | - | - | - | - | - | - | 1.25E-13 (7.41) |

[a] t-statistics in bracket. The statistics were summarized on 20,000 10-kb windows separated by 100 kb with each other.

**Supplementary Table S8. Migration weight estimated under different assumption of migration events.**

| Migration events | DROME→KAZA | DROME→RUS | DROME→IRAN | DROME→XJ_TLM | XJ_ML→XJ_ZGE |
|---|---|---|---|---|---|
| m=1 | 0.045362 | | | | |
| m=2 | 0.0998758 | 0.0839476 | | | |
| m=3 | 0.0799761 | 0.0691873 | 0.0397444 | | |
| m=4 | 0.0759546 | 0.0659031 | 0.0404412 | 0.00960696 | |
| m=5 | 0.0745299 | 0.0645908 | 0.0403732 | 0.00930027 | 0.114931 |

**Supplementary Table S9. Significant admixture (Z-score ≤ -2) by F3 tests.**

| Population Configuration | F3 Score | SD | Z-score |
|---|---|---|---|
| KAZA;IMG,drome | -0.00683953 | 1.56E-05 | -437.097 |
| KAZA;MG,drome | -0.00691661 | 1.61E-05 | -428.46 |
| KAZA;XJ,drome | -0.00529425 | 1.75E-05 | -302.969 |
| RUS;IMG,drome | -0.00297673 | 1.69E-05 | -176.393 |
| RUS;MG,drome | -0.00297742 | 1.75E-05 | -170.535 |
| RUS;XJ,drome | -0.00152741 | 1.91E-05 | -79.8165 |
| IRAN;IMG,drome | -0.000921691 | 1.96E-05 | -46.9863 |
| IRAN;MG,drome | -0.000826489 | 2.01E-05 | -41.0883 |
| KAZA;RUS,drome | -0.000511919 | 2.07E-05 | -24.7051 |

**Supplementary Table S10. Significant admixture (|Z-score| ≥ 2) by F4 tests.**

| Population Configuration [a] | F4 Score | SD | Z-score |
|---|---|---|---|
| IMG,XJ;wild,drome | 0.00152976 | 1.30E-05 | 117.919 |
| IMG,RUS;wild,drome | 0.00612666 | 1.79E-05 | 342.163 |
| IMG,KAZA;wild,drome | 0.00751604 | 1.66E-05 | 453.288 |
| IMG,IRAN;wild,drome | 0.00938352 | 2.30E-05 | 408.803 |
| wild,drome;MG,RUS | 0.00613598 | 1.85E-05 | 330.907 |
| wild,drome;MG,KAZA | 0.00752515 | 1.71E-05 | 441.032 |
| wild,drome;MG,IRAN | 0.00939432 | 2.31E-05 | 406.647 |
| wild,drome;XJ,MG | -0.0015388 | 1.31E-05 | -117.446 |
| wild,drome;XJ,RUS | 0.00459725 | 1.98E-05 | 232.171 |
| wild,drome;XJ,KAZA | 0.00598652 | 1.84E-05 | 325.54 |
| wild,drome;XJ,IRAN | 0.0078561 | 2.48E-05 | 317.417 |
| wild,drome;RUS,KAZA | 0.00138675 | 2.04E-05 | 67.82 |
| wild,drome;RUS,IRAN | 0.00325979 | 2.84E-05 | 114.693 |
| wild,drome;KAZA,IRAN | 0.00187257 | 2.69E-05 | 69.5758 |

[a] Configuration (Y, Z; wild, drome) is considered.

**Supplementary Table S11. Additional full length mtDNA sequences used for reconstructing the phylogenetic tree.**

| GenBank accession | Species | Sequence length (bp) | Origin |
|---|---|---|---|
| NC_009628 | Camelus bactrianus | 16659 | Inner Mongolia, China |
| KX554925 | Camelus bactrianus | 16659 | Iran |
| KX554926 | Camelus bactrianus | 16856 | Iran |
| KX554927 | Camelus bactrianus | 16669 | Iran |
| KX554928 | Camelus bactrianus | 16659 | Iran |
| KX554929 | Camelus bactrianus | 16659 | Iran |
| KX554930 | Camelus bactrianus | 16659 | Iran |
| AP003423 | Camelus bactrianus | 16663 | Unknown |
| EF507798 | Camelus bactrianus | 16659 | Umnugobi, Mongolia |
| EF507799 | Camelus bactrianus | 16667 | Gobi-Altai, Mongolia |
| EF212037 | Camelus bactrianus | 16659 | Inner Mongolia, China |
| MH109991 | Camelus bactrianus | 15432 | Iran |
| MH109997 | Camelus bactrianus | 15435 | Iran |
| NC_009629 | Camelus ferus | 16680 | Gobi area, Mongolia |
| EF212038 | Camelus ferus | 16680 | Gobi-Altai, Mongolia |
| EF507801 | Camelus ferus | 16651 | Gobi-Altai, Mongolia |
| EF507800 | Camelus ferus | 16670 | Gobi-Altai, Mongolia |
| NC_009849 | Camelus dromedarius | 16643 | United Arab Emirates: Dubai |
| KX554931 | Camelus dromedarius | 16643 | Iran |
| KX554932 | Camelus dromedarius | 16642 | Iran |
| KX554933 | Camelus dromedarius | 16622 | Iran |
| KX554934 | Camelus dromedarius | 16642 | Iran |
| KU605072 | Camelus dromedarius | 16379 | Qatar, Jordan border |
| KU605073 | Camelus dromedarius | 16379 | Saudi Arabia |
| KU605074 | Camelus dromedarius | 16379 | Saudi Arabia |
| KU605075 | Camelus dromedarius | 16379 | Saudi Arabia |
| KU605076 | Camelus dromedarius | 16379 | Unknown |
| KU605077 | Camelus dromedarius | 16379 | United Arab Emirates: Dubai |
| KU605078 | Camelus dromedarius | 16375 | Kenya |
| KU605079 | Camelus dromedarius | 16379 | Sudan |
| KU605080 | Camelus dromedarius | 16379 | Pakistan |
| JN632608 | Camelus dromedarius | 16665 | Morocco |

| | | | |
|---|---|---|---|
| EU159113 | Camelus dromedarius | 16643 | United Arab Emirates: Dubai |
| MH109998 | Camelus dromedarius | 15432 | Ardabili, Iran |
| MH109999 | Camelus dromedarius | 15432 | Ardabili, Iran |
| MH110000 | Camelus dromedarius | 15431 | Ardabili, Iran |
| MH110003 | Camelus dromedarius | 15434 | Ardabili, Iran |
| MH110004 | Camelus dromedarius | 15431 | Ardabili, Iran |
| MH110005 | Camelus dromedarius | 15432 | Ardabili, Iran |

**Supplementary Table S12. Nucleotide diversity of mtDNAs.**

| Population | Sample size | Nucleotide diversity |
|---|---|---|
| Drome | 4 | 0.00734 |
| IRAN | 6 | 0.00160 |
| XJ | 15 | 0.00143 |
| MG | 28 | 0.00143 |
| IMG | 30 | 0.00126 |
| Wild | 19 | 0.00107 |
| RUS | 10 | 0.01073 |
| RUS (removing introgression)[a] | 9 | 0.001 |
| QH | 10 | 0.00097 |
| KAZA | 6 | 0.00938 |
| KAZA (removing introgression)[b] | 5 | 0.00044 |

[a] SAMN06759189 is clustered with dromedaries.

[b] SAMN06759200 is clustered with dromedaries.

**Supplementary Table S13. G-PhoCS setup parameters.**

| Model | Parameter | Value |
|---|---|---|
| General settings | num-loci | 10000 |
| | burn-in | 100000 |
| | mcmc-iterations | 500000 |
| | mcmc-sample-skip | 10 |
| | locus-mut-rate | CONST |
| | find-finetunes | TRUE |
| | find-finetunes-num-steps | 100 |
| | find-finetunes-samples-per-step | 100 |
| | tau-theta-print | 10000 |
| | prior for all $\theta$ parameters | $\Gamma(\alpha = 1, \beta = 10000)$ |
| No migration | prior for $\tau$ A: (MG, KAZA) | $\Gamma(\alpha = 1, \beta = 100000)$ |
| | prior for $\tau$ B: (A, IRAN) | $\Gamma(\alpha = 1, \beta = 100000)$ |
| | prior for $\tau$ C: (B, wild) | $\Gamma(\alpha = 1, \beta = 10000)$ |
| | prior for $\tau$ root: (C, drome) | $\Gamma(\alpha = 1, \beta = 1000)$ |
| Migration | mig-rate-print | 0.1 |
| | prior for all m parameters | $\Gamma(\alpha = 0.002, \beta = 0.00001)$ |
| | migration bands | drome->IRAN, ghost->KAZA |
| | prior for $\tau$ A: (MG, KAZA) | $\Gamma(\alpha = 1, \beta = 100000)$ |
| | prior for $\tau$ B: (A, IRAN) | $\Gamma(\alpha = 1, \beta = 100000)$ |
| | prior for $\tau$ C: (B, wild) | $\Gamma(\alpha = 1, \beta = 10000)$ |
| | prior for $\tau$ X: (drome, ghost) | $\Gamma(\alpha = 1, \beta = 1000)$ |
| | prior for $\tau$ root: (C, X) | $\Gamma(\alpha = 1, \beta = 1000)$ |

**Supplementary Table S14. Parameter estimates of G-PhoCS summarized by Tracer.**

| Parameter [a] | No migration | | | Migration | | |
|---|---|---|---|---|---|---|
| | Mean | Stdev | 95% HPD interval [b] | Mean | Stdev | 95% HPD interval |
| θ MG | 0.73 | 0.75 | [4.60E-3, 2.26] | 15.46 | 3.15 | [9.70, 21.73] |
| θ KAZA | 2.79 | 1.62 | [0.21, 5.97] | 18.67 | 3.70 | [11.85, 26.11] |
| θ IRAN | 5.60 | 2.45 | [1.33, 10.46] | 36.93 | 5.62 | [26.03, 47.86] |
| θ wild | 1.55 | 0.28 | [1.02, 2.09] | 2.05 | 0.23 | [1.61, 2.49] |
| θ drome | 9.98 | 0.70 | [8.77, 11.47] | 8.98 | 0.64 | [7.87, 10.34] |
| θ ghost | - | - | - | 39.07 | 17.62 | [1.46, 64.69] |
| θ A | 3.33 | 1.67 | [0.60, 6.70] | 4.05 | 1.96 | [0.68, 7.98] |
| θ B | 3.12 | 0.51 | [2.17, 4.11] | 2.04E-2 | 1.54E-2 | [2.89E-3, 0.05] |
| θ C | 16.17 | 0.89 | [14.40, 17.91] | 13.33 | 0.70 | [12.00, 14.72] |
| θ X | - | - | - | 2.28 | 1.66 | [0.13, 5.41] |
| θ root | 30.44 | 0.68 | [29.13, 31.8] | 31.60 | 0.85 | [30.00, 33.33] |
| τ A | 5.10E-3 | 3.95E-3 | [1.00E-5,1.28E-2] | 0.46 | 1.99E-2 | [0.42, 0.50] |
| τ B | 7.07E-3 | 4.54E-3 | [1.70E-4,1.55E-2] | 0.46 | 1.98E-2 | [0.42, 0.50] |
| τ C | 0.34 | 5.35E-2 | [0.24, 0.45] | 0.46 | 0.02 | [0.42, 0.50] |
| τ X | - | - | - | 3.53 | 0.15 | [3.21, 3.82] |
| τ root | 3.96 | 0.14 | [3.67, 4.23] | 3.53 | 0.15 | [3.23, 3.84] |
| m drome->IRAN | - | - | - | 22.25 | 0.99 | [20.32, 24.20] |
| m ghost->KAZA | - | - | - | 21.70 | 0.96 | [19.82, 23.61] |

[a] θ and τ are scaled by $10^4$, m are scaled by 0.1.

[b] Highest posterior density interval. The HPD is a credible set that contains 95% of the sampled values.
